# Monitoring extracellular ion and metabolite dynamics with recombinant nanobody-fused biosensors

**DOI:** 10.1101/2022.04.12.488002

**Authors:** Sandra Burgstaller, Teresa R. Wagner, Helmut Bischof, Sarah Bueckle, Aman Padamsey, Desiree I. Frecot, Philipp D. Kaiser, David Skrabak, Roland Malli, Robert Lukowski, Ulrich Rothbauer

## Abstract

The tumor microenvironment (TME) consists of different cell types that secrete proteins and also control the extracellular concentration of ions and metabolites. Changes in these intra-tumoral analytes and conditions, including K^+^, glucose, and pH, have been described to alter the metabolic activity of cancer cells, promote tumor cell growth, and impair anti-tumor immunity. However, the mechanisms regulating ion and metabolite levels and their effects on certain characteristics of the TME are still poorly understood. Therefore, accurate determination and visualization of analyte or state changes in real time within the TME is desired.

In this study, we genetically combined FRET-based fluorescent biosensors with nanobodies (Nbs) and used them for targeted visualization and monitoring of extracellular changes in K^+^, pH, and glucose on cell surfaces. We demonstrated that these recombinant biosensors quantitatively visualize extracellular K^+^ alterations on multiple cancer and non-cancer cell lines and primary neurons. By implementing a HER2 specific Nb, we generated K^+^ and pH sensors, which retain their functionality and specifically stained HER2 positive breast cancer cells. Based on the successful technical development of several Nb-biosensor combinations, we anticipate that this approach can be easily extended to design other targeted biosensors. Such versatile probes will open new possibilities for the reliable study of extracellular analytes in advanced 3D cell models or even *in vivo* systems.

## INTRODUCTION

The tumor microenvironment (TME) represents a highly specialized niche, where tumor-associated-stromal cells, immune cells, blood-and lymphatic vessels create an oncogenic milieu featuring nutrients, growth factors, cytokines and abnormal alterations of intra-and extracellular ion and metabolite levels (Anderson and Simon, 2020). Cancer cell metabolism and proliferation are heavily influenced by these microenvironmental factors and their interaction (Costanza et al., 2019; Huntington et al., 2022; Pedersen et al., 2017; Soroceanu et al., 1999). In this context, the Warburg effect describes that cancer cells prefer glycolysis to oxidative phosphorylation despite the presence of molecular oxygen (O_2_) (Vander Heiden et al., 2009). This metabolic switch, also referred to as aerobic glycolysis, yields lactate production and subsequent secretion, thereby significantly acidifying the TME towards a pH of 6.5 or lower (Gallagher et al., 2008; Vander Heiden *et al*., 2009). This in turn affects cell and tissue organization. For example, acidic pH is known to trigger extracellular matrix (ECM) restructuring, leading to loss of ECM integrity (Busco et al., 2010; Webb et al., 2011) which facilitates the spread of cancer and tumor dissemination (Frantz et al., 2007). Associated with this phenomenon, the altered metabolism of cancer cells provides increased glucose uptake to meet the increased energy demand during proliferation (Han et al., 2015; Yang et al., 2013). Hence, the extracellular glucose concentration ([GLU]_ex_) represents a growth-determining factor (Cao et al., 2007; Xu et al., 2015). Moreover, the TME not only promotes cancer growth by metabolically reprogramming, but also by modulating responses of the immune system (Eil et al., 2016). As a result of improper vascularization, solid tumor growth is associated with necrotic cell death within the tumor core. During necrosis, high intracellular K^+^ levels are released into the TME, affecting the function of effector T-cells, ultimately causing cancer cells to escape the immune system (Eil *et al*., 2016). In summary, the interplay of such extracellular changes causes the development of a hostile environment that is presumed to impair the efficacy of chemotherapeutic agents (Jähde et al., 1990; Thews et al., 2011; Yang et al., 2020). Considering this, real-time monitoring of ionic and metabolic changes in the extracellular milieu could broaden our understanding of the bidirectional crosstalk between tumor cells and stromal cells. By maintaining a cellular resolution of these intra-tumoral signaling factors such strategies might also help to unravel unknown mechanisms that promote stromal cell recruitment by the tumor, cancer metabolism and cell malignancy.

Foerster resonance energy transfer (FRET)-based biosensors are powerful tools for measuring ions and analytes at the cellular level (Bischof et al., 2019; Burgstaller et al., 2022; Depaoli et al., 2019). These biosensors usually consist of two fluorescent proteins acting as FRET donor and acceptor, respectively, linked by an analyte-binding domain (Depaoli *et al*., 2019). The design of FRET-based biosensors enables reporting of changes in the analyte by increasing or decreasing FRET efficiency, a process that is fast, highly dynamic, and reversible (Depaoli *et al*., 2019). Most currently applied FRET-based sensors visualize intracellular analyte fluctuations. However, this requires genetic cell manipulation either by transient or stable transfection of the respective FRET biosensor, which limits their application due to drawbacks such as low transfection rates of primary cells or alteration of cell metabolic activities (Fiszer-Kierzkowska et al., 2011; Jacobsen et al., 2009; Mello de Queiroz et al., 2012). In contrast, FRET-based biosensors have also been applied as recombinant purified sensors to measure extracellular analytes (Bischof et al., 2017; Burgstaller et al., 2021; Namiki et al., 2007; Whitfield et al., 2015; Zhang et al., 2018). For cellular immobilization, these biosensors were further engineered using non-covalent biotin (strept/trapt) avidin interaction motifs; however, these approaches also rely on genetic manipulations of target cells or unspecific biotinylation of the cell surface (Burgstaller *et al*., 2021; Namiki *et al*., 2007; Whitfield *et al*., 2015; Zhang *et al*., 2018).

A more precise targeting specificity can be achieved by using antibodies or fragments thereof (Hamers-Casterman et al., 1993). Binding molecules derived from heavy chain only antibodies of camelids, termed VHHs or nanobodies (Nbs), have proven to be reliable tools for many applications in biomedical research, diagnostics, or even therapy (Liu et al., 2021; Muyldermans, 2021; Wagner and Rothbauer, 2021). Nbs are characterized by small size, antibody-like affinities and specificities, low off-target accumulation, high stability and good solubility (Muyldermans, 2013). Their unique properties and ease of genetic and/or chemical functionalization offer significant advantages over conventional antibodies. Recently, Nbs specific for GFP or RFP which were genetically fused with different variants of the fluorescent Ca^2+^ sensor GECO1.2 (Zhao et al., 2011) have been used to measure physiological Ca^2+^ changes in living cells after extrinsic stimulation. Similarly, the green fluorescent pH sensor super-ecliptic pHluorin (SEpHluorin) (Sankaranarayanan et al., 2000) or the red fluorescent pH sensor pHuji (Shen et al., 2014) and the excitation ratiometric ATP/ADP sensor Perceval-HR (Tantama et al., 2013) were combined with these Nbs to specifically measure pH shifts or ATP/ADP losses after inhibition of glycolysis and oxidative phosphorylation at distinct cellular compartments within live cells (Prole and Taylor, 2019).

In this study, we exploit the potential of Nbs to immobilize functional biosensors as recombinant proteins on the extracellular surface. Therefore, we developed biosensor fusion constructs using either a peptide tag-specific Nb (SPOT-Nb) (Braun et al., 2016) as a broadly applicable generic binding molecule, or a HER2-specific Nb (2Rs15d) (Vaneycken et al., 2011) targeting an endogenous surface protein in combination with GEPII 1.0 (Bischof *et al*., 2017), pH-Lemon (Burgstaller et al., 2019), or FLII12Pglu-700µδ6 (further referred to as FLII) (Takanaga et al., 2008) to measure extracellular K^+^, pH, and glucose changes near the cell surface. Our results showed that these sensors can be successfully immobilized on the cell surface and retain their full functionality. Most importantly, the Nb-fused biosensors enabled spatially resolved physiologically relevant FRET-based or fluorescent measurements of extracellular changes in all analytes tested over an extended period of time. From our findings, we propose that this versatile approach opens new opportunities to study important metabolic activities at the interface of cells and the extracellular matrix (ECM) in advanced experimental settings including 3D organoids or even *in vivo* models.

## RESULTS

To measure changes in K^+^, pH, and glucose as important TME-associated analytes in the extracellular space, three FRET biosensors were used. GEPII 1.0 represents a highly specific indicator for K^+^, which is based on a conformational rearrangement mediated by the K^+^ binding protein Kbp (**Figure 1A**) (Bischof *et al*., 2017). pH-Lemon, a pH sensor, which is based on the intrinsic pH sensitivity and insensitivity of EYFP and mTurquoise2, respectively (**Figure 1B**) (Burgstaller *et al*., 2019), and FLII, a widely used glucose indicator, which permits glucose sensing by conformational changes of the glucose binding domain, MglB, thereby increasing FRET (**Figure 1C**) (Takanaga *et al*., 2008).

**Figure 1.**
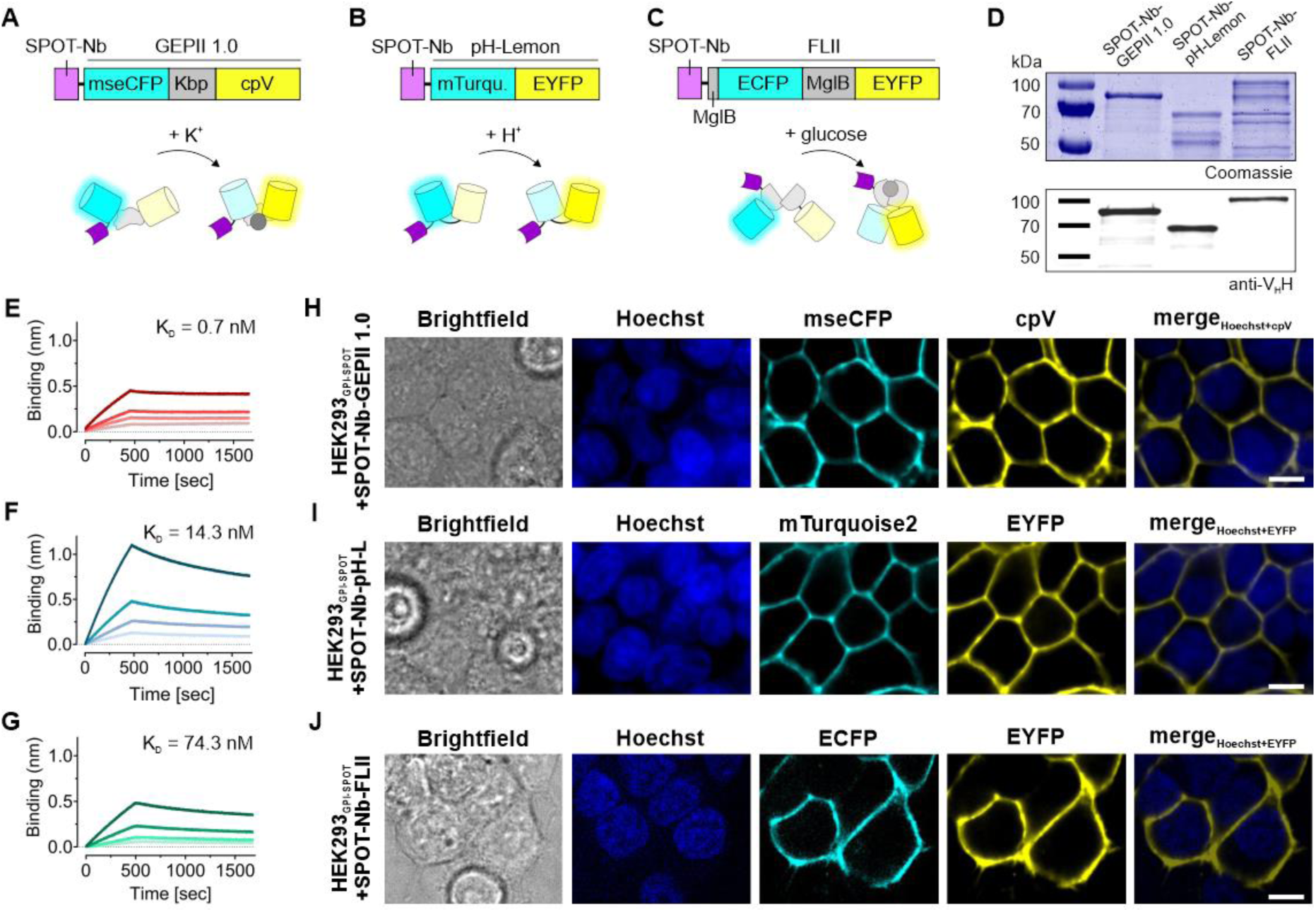
Fluorescent biosensors fused to the SPOT-Nb specifically bind the SPOT-tag. Schematic illustration of (**A**) the SPOT-Nb-GEPII 1.0 consisting of the Nb (magenta), the FRET donor (mseCFP, cyan), the potassium binding domain (Kbp, grey) and the FRET acceptor (cpV, yellow), responding to K^+^ alterations by conformational rearrangement of Kbp, (**B**) the SPOT-Nb-pH-Lemon consisting of the Nb (magenta) fused to the pH stable mTurquoise2 (cyan), and the pH sensitive EYFP (yellow), responding to pH alterations due to quenching of the EYFP fluorescence, (**C**) the SPOT-Nb-FLII comprising the Nb (magenta) domain, a FRET donor (ECFP, cyan), the separated glucose binding domains (MglB, grey) and the FRET acceptor (EYFP, yellow), responding to glucose alterations by conformational rearrangement of the MglB domains. (**D**) Recombinant expression and purification of the Nb-fused biosensors using immobilized metal ion chromatography (IMAC) and size exclusion (SEC). Coomassie-stained SDS-PAGE of 1 µg (upper panel) and immunoblot analysis using anti-V_H_H antibody (lower panel) of purified SPOT-Nb-GEPII 1.0, SPOT-Nb-pH-Lemon and SPOT-Nb-FLII proteins are shown. (**E - G**) For biolayer interferometry (BLI)-based affinity measurements, biotinylated SPOT peptide was immobilized on streptavidin biosensors. Kinetic measurements were performed using four concentrations of purified Nb-fused biosensors ranging from 6.25 – 100 nM. Sensograms of SPOT-Nb-GEPII 1.0 (**E**), SPOT-Nb-pH Lemon (**F**) and SPOT-Nb-FLII (**G**) are shown. (**H - J**) Representative confocal microscopy images of live HEK293 cells expressing GPI-anchored SPOT-tag (GPI-SPOT) on the plasma membrane upon incubation with SPOT-Nb-GEPII 1.0 (**H**); SPOT-Nb-pH Lemon (**I**) and SPOT-Nb-FLII (**J**). Shown are from left to right: Brightfield, Hoechst, and the fluorescent signals of the respective FPs as indicated. Scale bar 10 µm.

To generate the Nb-fused biosensors, the SPOT-Nb was fused N-terminally to the different FRET pairs (**Figures 1A-1C**) (Braun *et al*., 2016). This well-established Nb binds the SPOT peptide (SPOT-tag), with high specificity and affinity and provides optimal properties for validating the applicability of Nb-mediated immobilization of biosensors on the cell surface. In a first step, all Nb-fused biosensors were cloned with a C-terminal His_6_-tag, expressed in *Escherichia coli* (*E.coli*) and purified using immobilized metal ion affinity chromatography (IMAC) followed by size exclusion chromatography (SEC). SDS-PAGE analysis from the last purification step showed that all three sensor constructs are expressed and yielded at the expected molecular weight as full-length sensor fusion protein of SPOT-Nb-GEPII 1.0 (86 kDa), SPOT-Nb-pH-Lemon (71 kDa) and SPOT-Nb-FLII (103 kDa) (**Figure 1D**). However, for SPOT-Nb-pH-Lemon and SPOT-Nb-FLII additional bands referring to smaller proteins were detected. Immunoblot analysis using an anti-V_H_H antibody revealed that the smaller proteins were lacking the SPOT-Nb moiety (**Figure 1D**). These findings indicated that the chimeric SPOT-Nb-pH-Lemon and SPOT-Nb-FLII constructs are sensitive to degradation that occurs throughout the expression or purification process. However, since the resulting protein fragments could not bind the SPOT-tag, we concluded that they should not interfere with subsequent measurements based on Nb binding and sensor functionality. Next, we investigated whether the SPOT-Nb (∼15 kDa) fused to the large sensor proteins (∼ 52 - 88 kDa) exhibits still full binding capacity. To this end, we determined the binding affinities of the Nb-fused biosensors to the isolated SPOT-peptide by biolayer interferometry. Strong binding of the Nb-fused biosensors to the SPOT-tag with dissociation rate constants (K_D_) in the low nanomolar range of ∼ 0.7 nM and ∼14.3 nM for SPOT-Nb-GEPII 1.0 (**Figure 1E and Table S1**) and SPOT-Nb-pH-Lemon (**Figure 1F and Table S1**), respectively were determined. These affinities are comparable to those previously measured for the SPOT-Nb alone (Braun *et al*., 2016). However, a slightly higher K_D_ of ∼ 74.3 nM was determined for the SPOT-Nb-FLII (**Figure 1G and Table S1**) which could be due to a steric hindrance caused by the large biosensor moiety of ∼ 88 kDa.

To test the binding properties in a more relevant setting, we examined the ability of the purified SPOT-Nb-fused biosensors to bind to GPI-anchored SPOT tags (GPI-SPOT) on the surface of HEK293 cells. Therefore, HEK293 either transiently transfected with a GPI-SPOT expression construct or left untreated (HEK293 wildtype (WT)), were incubated with the SPOT- Nb-biosensor fusion constructs (**Figuress 1H-1J** and **S1A**), or a fluorescently labeled SPOT-Nb (SPOT-Nb_ATTO488_) with a small ATTO488 fluorophore (∼1 kDa) as a positive control (**Figure S1B**). Live-cell fluorescent imaging resulted in strong fluorescence signals exclusively localized at the plasma membrane of HEK293 cells expressing the GPI-SPOT (**Figures 1H-1J**). The obtained signal intensities and localization were comparable to the respective SPOT-Nb_ATTO488_ staining (**Figure S1B**). In contrast, no fluorescence signals were detected on non-transfected HEK293 cells (**Figure S1A**) or upon incubation of cells expressing GPI-SPOT with recombinant purified GEPII 1.0 lacking the SPOT-Nb (**Figure S1C**). From these findings, we concluded that fusion of the SPOT-Nb to the sensors did not affect the binding properties of the Nb and that all tested SPOT-Nb-fused biosensors are suitable to identify cells presenting the SPOT-tag on the extracellular surface of their plasma membrane.

Next, we investigated the stability of binding and potential internalization of the SPOT-Nb biosensors over time, as both could limit the reliable and sustained measurement of extracellular analytes. Thus, GPI-SPOT expressing HEK293 cells were incubated with SPOT-Nb-GEPII 1.0 and SPOT-Nb-pH-Lemon and time-lapse imaging of the fluorescence signals of the biosensors was performed for 4.5 hours. For both Nb-fused biosensors, the corresponding fluorescent images showed that signals were retained specifically at the plasma membrane. Endocytotic vesicles containing fluorescence signals were rarely observed (**Figures S2A** and **S2B**). To follow these processes for an even longer period, we continued and additionally re-investigated the cells after 48 hours of initial immobilization. By this, we observed that all tested Nb-fused biosensors were nearly undetectable 48 hours after immobilization, possibly due to disassociation or degradation. However, re-loading with SPOT-Nb-GEPII 1.0 resulted in a similar membrane staining pattern compared to earlier time points (**Figure S3**). Overall, these results showed that the biosensor constructs fused with SPOT-Nb are capable of transiently and specifically targeting the plasma membrane over an extended period of time. We further concluded that these sensors can be used for long-term experiments, although cell re-staining may be required.

So far, our results have shown that the biosensors fused with SPOT-Nb can specifically recognize the antigen expressed on the surface of living cells. Hence, the next question was whether the immobilized biosensors are capable of dynamically detecting corresponding changes in extracellular analytes with an appropriate signal-to-noise ratio and sensitivity. First, we examined the fluorescence emission signals of mseCFP and FRET of SPOT-Nb-GEPII 1.0 interacting with GPI-SPOT at the surface of HEK293 cells in response to the administration of buffers with different K^+^ concentrations ([K^+^]). Before the analysis, we confirmed the correct localization of the Nb-fused biosensor by fluorescent live-cell imaging revealing specific staining of the plasma membrane (**Figures S4A and S4B**). Subsequently, the cells were perfused with buffers comprising increasing [K^+^] ranging from 0 – 100 mM, and FRET signals were continuously visualized (**Figure 2A**). In line with the [K^+^]-dependent FRET signals obtained (**Figure 2A**), the fluorescence emission signals of the single FPs displayed a ratiometric behavior with decreasing mseCFP and increasing FRET fluorescence emissions (**Figure 2B**). The calculated half-maximal concentration (EC_50_) of 8.6 mM (**Figure 2C**), indicated that SPOT-Nb-GEPII 1.0 covers a (patho-) physiologically relevant range of extracellular [K^+^] ([K^+^_ex_]) (Eil *et al*., 2016). Next, we investigated the sensitivity of the immobilized SPOT-Nb-pH Lemon. For this purpose, cells were exposed to different extracellular pH values (pH 5 – 9). Upon separate excitation of mTurquoise2 and EYFP, as previously reported (Burgstaller *et al*., 2019), SPOT-Nb-pH Lemon showed ratiometric changes dependent on extracellular pH (pH_ex_) (**Figures 2D and S4C**). Based on the sensor design, SPOT-Nb-pH-Lemon repeatedly responded to decreasing pH_ex_ with increasing mTurquoise2 fluorescence due to the pH stability of FP, which as a FRET donor, however, is affected by the attached pH-sensitive EYFP. As expected, decreasing EYFP fluorescence could be observed upon acidification, caused by the protonation of the fluorophore and, subsequently, quenching of the FP (**Figures 2E and S4D**). Considering the calculated pKa value of 7.0 (**Figure 2F**), which corresponds well with the previously reported pH-sensitivity of the sensor (Burgstaller *et al*., 2019; Burgstaller *et al*., 2021), we concluded that immobilized SPOT-Nb-pH Lemon is suitable to detect physiologically relevant changes in pH_ex_ (Gallagher *et al*., 2008). In contrast to Nb-fused GEPII 1.0 and pH-Lemon, the FLII biosensor exhibits a more complex biosensor design as it relies on the formation of a functional glucose binding domain based on two split MglB fragments (Deuschle et al., 2005; Takanaga *et al*., 2008). However, as we inserted a long flexible linker between the Nb and the first MglB fragment we hypothesized that this design would confer sufficient steric flexibility for glucose binding and FRET signal generation. Hence, the functionality of SPOT-Nb-FLII to detect alterations of the extracellular glucose concentration ([GLU]_ex_) was tested upon cellular immobilization on HEK293 GPI-SPOT cells (**Figures S4E and S4F**). Cells were perfused with buffers comprising increasing [Glu]_ex_ ranging from 0 – 100 mM (**Figure 2G**). Continuous measurement of the FRET ratio signals revealed a distinct, ratiometric response of the immobilized Nb-fused biosensor to [GLU]_ex_ alterations in the low mM range (**Figures 2G and 2H**) with an estimated EC_50_ of 0.9 mM (**Figure 2I**). These findings demonstrated that all SPOT-Nb fused biosensors retain their functionality and report changes in extracellular analytes and conditions in a physiologically relevant range. Apparently, the N-terminal Nb-binding moiety does not negatively affect functional biosensor conformation, which may be due to the presence of a long flexible linker between the two functional domains.

**Figure 2.**
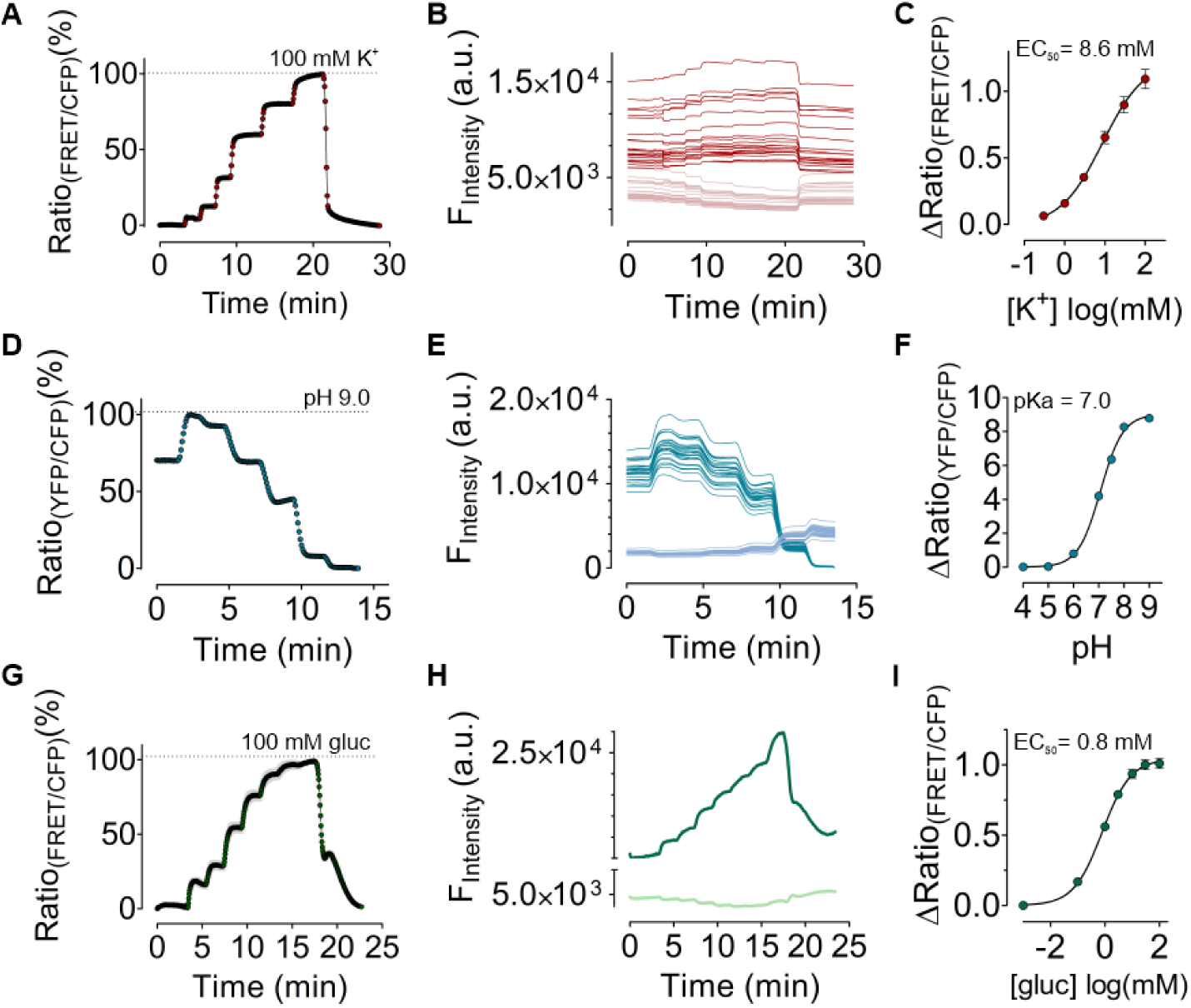
Biosensors immobilized on the plasma membrane respond to K^+^, pH and glucose alterations. (**A**) Response of SPOT-Nb-GEPII 1.0 immobilized on HEK293 cells expressing GPI-SPOT upon perfusion of buffers with different [K^+^] over time. Shown is the mean ± SD of one representative measurement of multiple cells as shown in (**B**). (**B**) Single cell traces of cpV (red) and mseCFP (pink) of immobilized SPOT-Nb-GEPII 1.0 in response to buffers with different [K^+^] as shown in panel (**A**). (**C**) Dose response curve of SPOT-Nb-GEPII 1.0 with an EC_50_ of 8,6 mM (4,8 mM – 15,5 mM). Shown is the mean ± SEM of four measurements representing biological replicates. (**D**) Response of SPOT-Nb-pH-Lemon immobilized on HEK293 cells expressing GPI-SPOT upon perfusion of buffers with different pH over time. Shown is the mean ± SD of one representative measurement of multiple cells as shown in (**E**). (**E**) Single cell traces of mTurquoise2 (blue) and EYFP (petrol) of immobilized SPOT-Nb-pH-Lemon in response to buffers with different pH as shown in panel (**D**). (**F**) Dose response curve of SPOT-Nb-pH-Lemon with a pKa of 7.0 (7.0 – 7.1) Shown is the mean ± SEM of three measurements representing biological replicates. (**G**) Ratiometric response of SPOT-Nb- FLII immobilized on HEK293 cells expressing GPI-SPOT upon perfusion of buffers with different glucose levels over time. Shown is the mean ± SD of one representative measurement of multiple cells. (**H**) Single cell traces of ECFP (light green) and EYFP (dark green) immobilized SPOT-Nb- FLII in response to buffers with different pH as shown in panel (**G**). (**I**) Dose response curve of SPOT-Nb- FLII with an EC_50_ of 0,8 mM (0,6 mM – 1 mM). Shown is the mean ± SEM of three measurements representing biological replicates.

Considering that we have so far evaluated Nb-fused biosensors exclusively for the detection of externally induced [K^+^]_ex_, pH_ex_ and [GLU]_ex_ alterations, we next aimed to investigate the suitability of our approach for monitoring endogenously elicited signals. To demonstrate the relevance and feasibility of our approach, we analyzed the performance of the SPOT-Nb-GEPII 1.0 to detect changes in [K^+^]_ex_ using primary hippocampal mouse neurons. In a proof-of-concept approach, neurons transiently expressing the GPI-SPOT construct were incubated with glutamate that massively increases [K^+^]_ex_ (**Figure 3A**) (Burgstaller *et al*., 2021; Ehinger et al., 2021; Hösli et al., 1981). Immobilization of the SPOT-Nb-GEPII on primary hippocampal mouse neurons was verified by live-cell imaging as described (**Figure 3B**). In the following, we performed real time FRET analysis and monitored glutamate-mediated release of endogenous K^+^ to the extracellular space. First, we added pure buffer, followed by treating the cells with a glutamate bolus. Subsequently, a perfusion-mediated K^+^ wash out was performed, followed by the addition of 100 mM [K^+^]_ex_. The FRET ratio signals of SPOT-Nb-GEPII 1.0 remained virtually unaffected upon the addition of pure buffer, whereas injection of glutamate immediately increased FRET ratio signals, indicating glutamate-triggered K^+^ efflux from individual neurons (**Figures 3C and 3E**). Interestingly, the recorded FRET signals did not differ significantly under these conditions, either at the soma (**Figure 3D**) or at the dendrites (**Figure 3E**), suggesting K^+^ is either evenly released from different cellular compartments or that the changes in [K^+^]_ex_ are subsequently detected by sensor molecules immobilized on different parts of the neurons. To ensure sensor functionality and classify the glutamate-induced FRET signals, FRET generation and decline were assessed in response to a K^+^-free or a buffer containing 100 mM K^+^ (**Figures 3D and 3E**). Importantly, our observations are in line with the previously reported decline of intracellular FRET [K^+^]_i_ signals monitored by GEPII 1.0 in glutamate exposed primary cerebellar granule cells (Ehinger *et al*., 2021). Hence, this proof-of-concept study indicates that the SPOT-Nb-GEPII 1.0 is also functional to report and monitor (patho)physiological K^+^ efflux from neurons.

**Figure 3.**
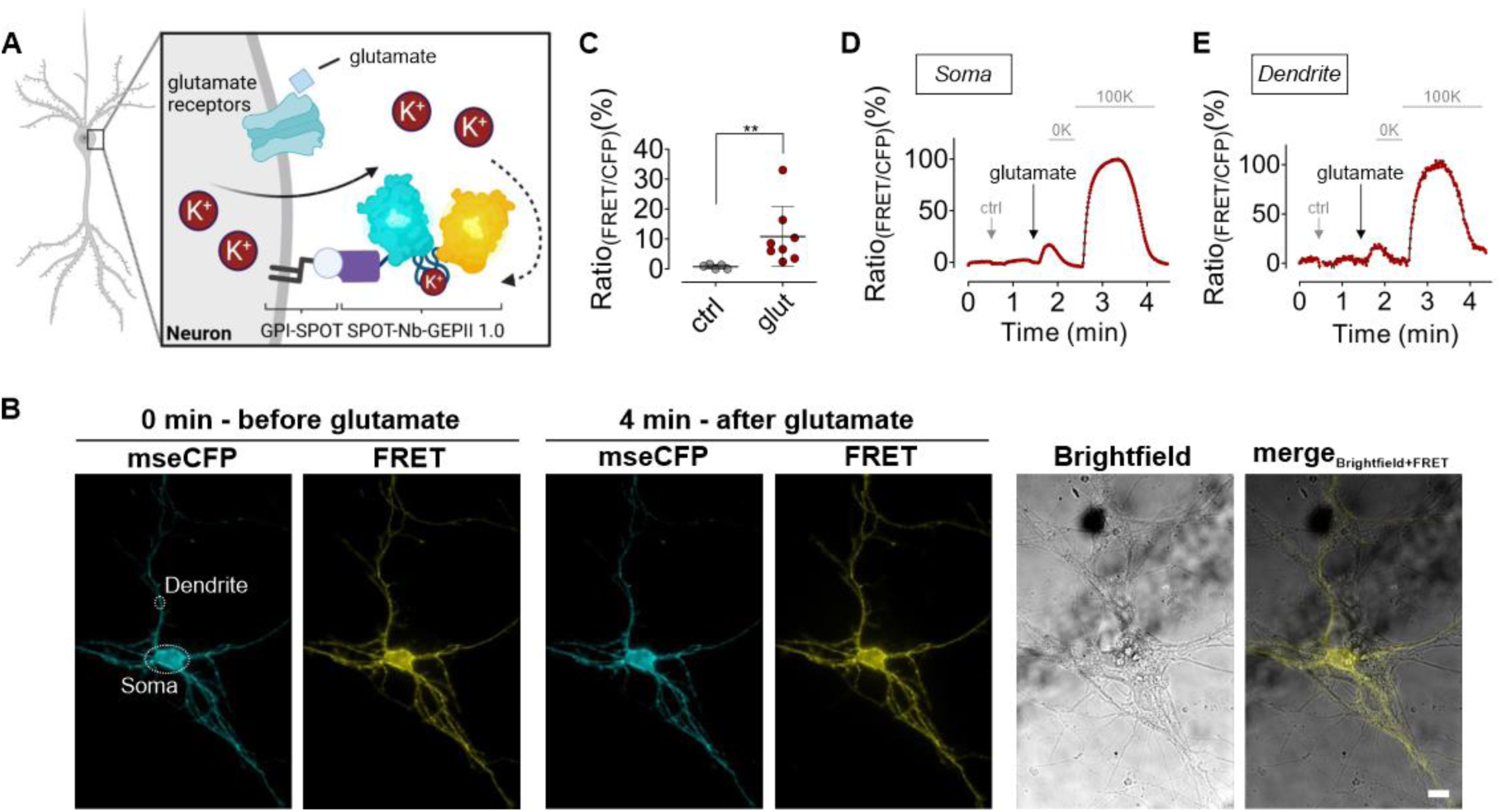
K^+^ sensors immobilized using SPOT-Nb can be used to track neuronal, glutamate-induced K^+^ efflux. (**A**) Schematic illustration of glutamate-induced K^+^ efflux from primary hippocampal mouse neurons. Glutamate (light blue) binding to glutamatergic receptors (glutamate receptors, light green) located in the plasma membrane causes K^+^ efflux, which can be monitored by SPOT-Nb-GEPII 1.0 immobilized on the surface of primary hippocampal mouse neurons expressing GPI-SPOT. (**B**) mseCFP and FRET widefield images of SPOT-Nb-GEPII 1.0 immobilized on the membrane of a primary hippocampal neuron before (0 min) and after (4 min) addition of glutamate, washing and re-addition of K^+^, as well as brightfield and a merge of brightfield and EYFP are shown. White circles indicate measurement region as shown in (**D**) and (**E**) at the soma and dendrites. Scale bar 10 µm. (**C**) Average (black lines) and single cell ratio changes in response to addition of a vehicle (ctrl, grey dots) and glutamate (red dots). Mann Whitney test, p = 0.0016. (**D,E**) Ratiometric response of SPOT-Nb-GEPII 1.0 upon injection of a vehicle control (1^st^ arrow, ctrl), injection of glutamate (2^nd^ arrow) and upon perfusion with K^+^-free buffer (0K) and subsequently with buffer containing 100 mM K^+^ (100K) measured at the soma (**D**) or at the dendrite (**E**) as indicated in (**B**).

Notably, all these results were obtained with SPOT-Nb fused biosensors, which require the expression of the SPOT-tag as a broadly applicable but artificial antigen on the plasma membrane. To further analyze whether this approach can also be used for endogenous cell surface epitopes, we selected an Nb (2Rs15d), which has been previously described to bind human epidermal growth factor receptor 2 (HER2) - a plasma membrane receptor widely expressed in breast cancer (Vaneycken *et al*., 2011). Following our original strategy, we designed and generated HER2-Nb-fused GEPII 1.0 or pH-Lemon biosensors in analogy to SPOT-Nb-based constructs. Bacterial expression yielded intact recombinant proteins showing only minor degradation as demonstrated by SDS-PAGE and immunoblotting (**Figure 4A**). We further performed live-cell imaging analysis on HEK293 cells transiently expressing human HER2. Cells staining with purified HER2-Nb-GEPII 1.0 (**Figure 4B**) and HER2-Nb-pH-Lemon (**Figure 4C**) resulted in prominent staining of the plasma membrane, while untransfected HEK293 cells remained unstained (**Figures 4B and 4C**). Additionally, incubation of HEK293 cells expressing HER2 with GEPII 1.0 lacking the Nb also showed no fluorescence signal (**Figure S5**). The imaging results indicated that both HER2-Nb fused biosensors can specifically recognize and properly bind their cell-surface target.

**Figure 4.**
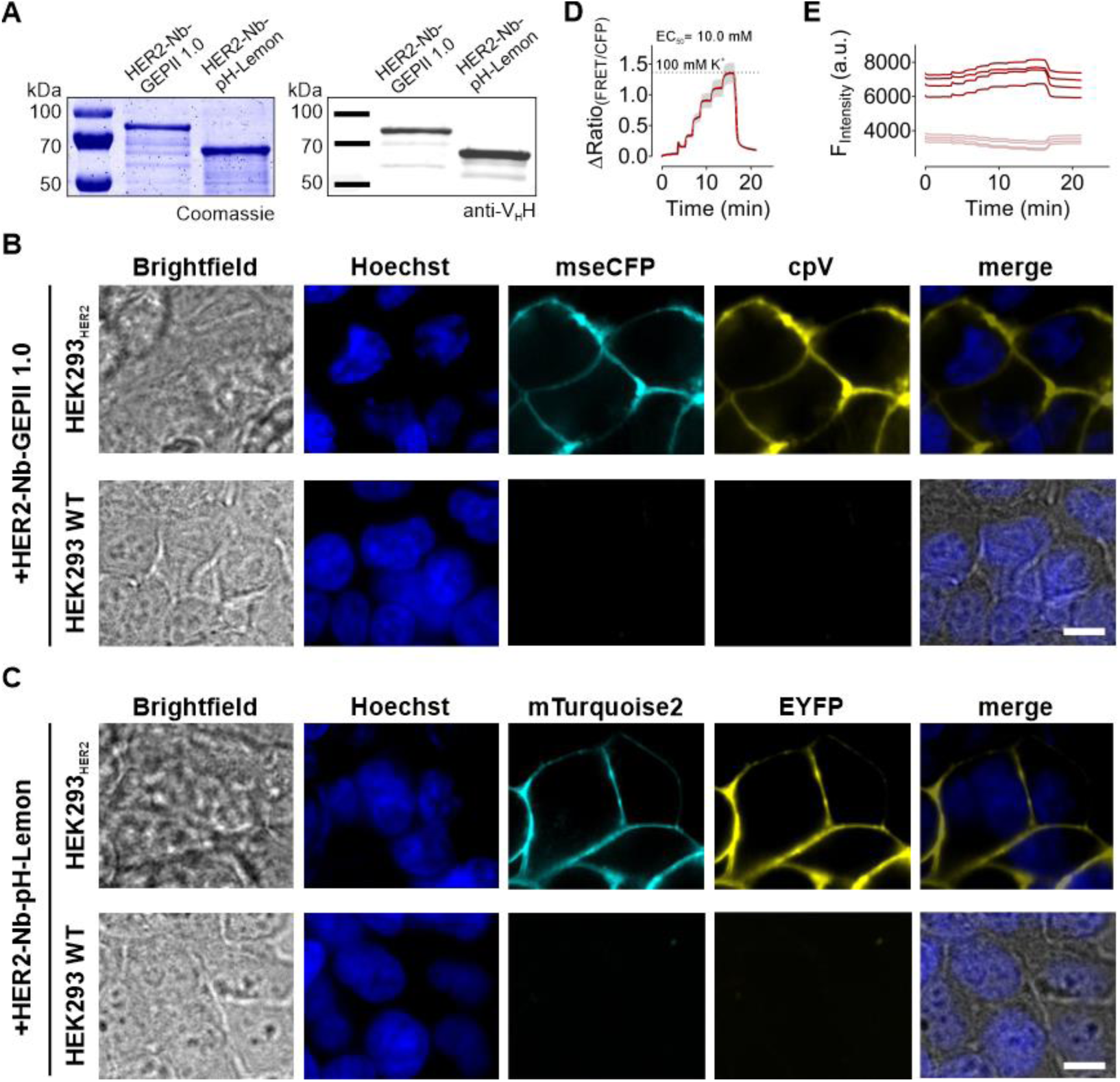
Biosensors fused to HER2-Nb precisely report [K^+^]_ex_ changes upon immobilization on HER2 expressing HEK293. (**A**) Coomassie-stained SDS PAGE of 1 µg protein (left panel) and immunoblot analysis using anti-VHH antibody (right panel) of purified HER2-Nb-GEPII 1.0 and HER2-Nb-pH-Lemon proteins are shown. (**B,C**) Representative confocal images of living HEK293 cells transiently overexpressing HER2 (upper row) or untransfected HEK293 cells (lower row) following incubation with HER2-Nb-GEPII 1.0 (**B**) or HER2-Nb-pH-Lemon (**C**) are shown. Scale bar 10 µm, n= 4 experiments representing biological replicates. (**D**) Response of HER2-Nb-GEPII 1.0 immobilized on HEK293 cell transiently overexpressing HER2 in response to buffers with different K^+^. Shown is a representative curve (mean ± SD from one representative measurement of multiple cells). (**E**) Respective single wavelength traces (FRET in red, mseCFP in pink) of the ratio curve as shown in (**D**) in response to K^+^ alterations.

To elucidate the functionality of the biosensors, we tested HER2-Nb-GEPII 1.0, as its functional principle is more complex due to the conformational change, compared to the fluorescence quenching of pH-Lemon. Perfusion of HER2 expressing HEK293 with increasing [K^+^]_ex_ (0 – 100 mM) revealed the functionality of HER2-Nb-GEPII 1.0 construct for detecting alterations in the [K^+^]_ex_ (**Figure 4D**). Similarly, HER2 Nb-fused GEPII 1.0 exhibited ratiometric behavior with increasing FRET and decreasing mseCFP fluorescence in response to increasing [K^+^]_ex_, revealing an EC_50_ value for K^+^ of 10 mM (**Figures 4D and 4E**) which is well in the range of SPOT-Nb-GEPII 1.0 (**Figure 2C**).

Having validated their functionality, we finally tested whether the recombinant HER2-Nb-fused biosensors can be immobilized on cells endogenously expressing HER2. Therefore, we utilized two breast cancer cell lines either positive (SkBr3) or negative (MCF7) for human HER2 (**Figure 5A**). Upon incubation with both Nb-fused biosensors, live-cell imaging showed a clear fluorescence staining of the cell surface for HER2-Nb-GEPII 1.0 and HER2-Nb-pH-Lemon for SkBr3 cells (**Figure 5B**) but not for MCF7 cells (**Figure 5C**). The HER2-Nb mediated specificity was further confirmed by the absence of any fluorescent staining of SkBr3 and MCF-7 cells upon incubation with GEPII 1.0 lacking the HER2-Nb (**Figure S5**). Additional time-lapse imaging showed a stable signal at the plasma membrane for 4.5 hours, despite some endocytosed vesicles (**Figures S6A and S6B**), which may indicate that cell viability is not affected by immobilization of the biosensors via the HER2-Nb. In line with these findings, an MTT-based assay showed that labeling of SkBr3 cells with HER2-Nb or HER2-Nb-GEPII 1.0 did not affect cell viabilities compared with control cells at 3, 6, 24, and 48 hours (**Figure 5D**). Based on these observations, we assume that due to stable immobilization in combination with low cell toxicity, the recombinant Nb-fused biosensors can also be used for more complex models such as 3 D cell models or patient-derived organoids.

**Figure 5.**
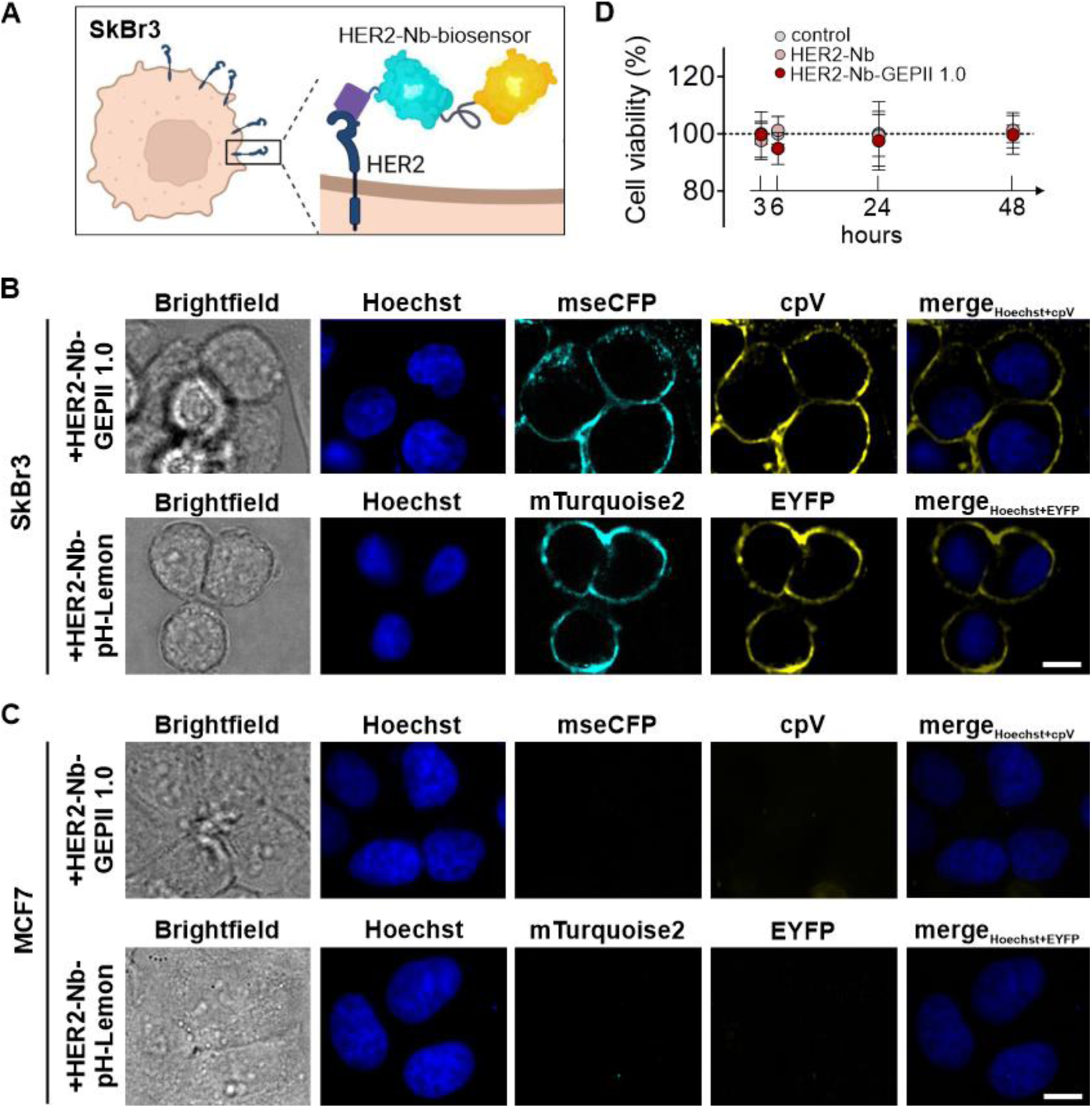
HER2-Nb-biosensors specifically label endogenous HER2 on HER2 positive breast cancer cells. (**A**) Schematic illustration of a HER2 positive SkBr3 cell endogenously expressing HER2 on the cell surface. Biosensors fused to HER2 can be bound to HER2 for immobilization on the plasma membrane. (**B**) Representative images of SkBr3 cells following incubation with HER2-Nb-GEPII 1.0 and HER2-Nb-pH-Lemon. Scale bar 10 µm, n= 4 experiments representing biological replicates. (**C**) Representative images of HER2 negative MCF7 breast cancer cells following incubation with HER2-Nb-GEPII 1.0 and HER2-Nb-pH-Lemon. Scale bar 10 µm, n= 4 experiments representing biological replicates. (**D**) Determination of cell viability of SkBr3 cells using MTT in response to vehicle (ctrl), unfused HER2-Nb and HER2-Nb-GEPII 1.0 after 3 h, 6 h, 24 h, and 48 h after immobilization. Not significant as determined using 1-way ANOVA, Dunn’s Multiple Comparison Test.

## DISCUSSION

Here, we provide evidence that recombinant Nb-fused FRET biosensors enable the visualization of dynamic analyte alterations in the extracellular compartment. Changes in ions and metabolites were visualized in a single-cell resolution and in real-time. Such approaches are relevant for optical mapping of pathophysiological relevant alterations of discrete analytes in extracellular fluids i.e., in the vasculature, lymphatic vessels, body cavities, synovial compartments, the cerebrospinal fluid as well as the TME. Moreover, the flexible combination of various, target-specific Nbs with different sensor molecules makes our toolbox approach versatile applicable.

The obtained results indicate that both - heterologously expressed and endogenous plasma membrane epitopes - are specifically targeted using different nanobodies such as SPOT-Nb (Braun *et al*., 2016) and HER2-Nb (Vaneycken *et al*., 2011). Importantly, immobilization of the Nb-fused FRET biosensors on their target side does not interfere with the conformational rearrangement as shown for the potassium sensor GEPII 1.0 (Bischof *et al*., 2017) and the glucose sensor FLII12Pglu700µd6 (Takanaga *et al*., 2008), nor the extinction of YFP fluorescence at low pH as shown for pH-Lemon (Burgstaller *et al*., 2019), which are necessary to produce accurate and reliable FRET signals, relying on different sensor principles. Although different in the mode of action, biosensors immobilized on the cell surface expressing the Nb targets remained fully functional for visualizing physiologically relevant changes in K^+^, pH, and glucose (Eil *et al*., 2016; Gallagher *et al*., 2008; Takanaga *et al*., 2008). The respective EC_50_ values obtained upon Nb immobilization were comparable to previously reported values (Burgstaller *et al*., 2021; Takanaga *et al*., 2008). *Vice versa*, the fusion of quite bulky biosensors to the much smaller Nb did not alter the affinity of the Nb. Accordingly, Nb-fused FRET biosensors displayed similar affinities in the low nanomolar range despite their significant increase in molecular weight compared to the unfused Nb (Braun *et al*., 2016; Vaneycken *et al*., 2011). Notably, for the generation of the Nb-fused biosensors, we chose a flexible design with long linker sequences that allow maximum steric freedom of the functional subunits. Although all Nb-fused biosensors could be functionally expressed in bacterial systems, they differed in their protein integrity and stability, which could be due to the different requirements needed for the correct formation of each functional subunit. In particular, the proper formation of essential disulfide bonds within the immunoglobulin fold of Nb requires the oxidative environment of the bacterial intermembrane space, whereas FPs such as CFP or YFP are preferentially expressed in the reducing environment of the cytoplasm (Feilmeier et al., 2000; Jain et al., 2001; Kunz et al., 2018). To improve expression conditions in the future, additional expression of chaperones such as disulfide isomerase (DsbC) should be considered (Olichon and Surrey, 2007).

By now, multiple FRET-based approaches were developed to monitor analytes in a single cell compartment or within the entire cells and thus are introduced as genetically encoded constructs (Depaoli *et al*., 2019). However, genetic manipulation of sensor-targeted cells bears the risk of low transfection efficiencies and/or metabolic alterations that might interfere with the readout of interest (Fiszer-Kierzkowska *et al*., 2011; Mello de Queiroz *et al*., 2012). Various strategies have already been described for the specific targeting of biosensors as recombinant proteins, ranging from the well-known biotin-avidin interaction (Burgstaller *et al*., 2022; Zhang *et al*., 2018) to insertion peptides such as pHLIPs (Weerakkody et al., 2013), which are integrated into the plasma membrane in response to low pH. Due to the acidic pH within the TME, this technique might be suitable to address this compartment in general. However, pHLIPs cannot be used to target specific proteins e.g. on cancer cells. In contrast, by using Nbs as the smallest intact antigen-binding fragments specific for cell surface markers, our approach focuses on their application to the specific study of extracellular signal mediators. The Nb-fused GEPII 1.0 sensor presented here could readily be used to study the effects of discrete plasma membrane K^+^ channels that facilitate the flux of K^+^ ions between the cytosol and the extracellular space. Such channels show cancer specific expression patterns and are present in a variety of solid tumors (Mohr et al., 2019c; Pardo and Stühmer, 2014; Steudel et al., 2017). Notably, recently agents modulating these channels have demonstrated antitumor efficacy (Mohr et al., 2019b; Mohr et al., 2020; Peruzzo and Szabo, 2019). Accordingly, our approach of a cell-specific immobilization of K^+^ biosensors might facilitate the screening and identification of such channel modulating agents. In another aspect, a recent analysis of genetically modified mouse models lacking, for instance, distinct Ca^2+^-activated K^+^ channels (KCa) revealed that this family of K^+^ channels is crucially involved in breast cancer development and its response to therapy (Mohr et al., 2019a; Steudel *et al*., 2017). However, the contribution of cancer-associated K^+^ channels to the TME and how their activity relates to functionally relevant changes in [K^+^]_ex_ is poorly understood (Burgstaller *et al*., 2022). Therefore, we anticipate that further development of our HER2-Nb-GEPII 1.0 for 3D cell models or *in vivo* applications in these models will lead to a more comprehensive understanding of the impact of this class of “onco-channels” (Huber, 2013). Previously, several studies suggested that the ionic composition and thus [H^+^]_ex_ and [K^+^]_ex_ play a role in the stromal cell recruitment at primary and metastatic tumor sites and in anti-tumor immunity (Comes et al., 2015; Huang et al., 2016; Som et al., 2016). By using combinations of the Nb-fused FRET biosensors developed here, the interplay of extracellular ions, metabolites, and conditions in response to various stimuli, such as cancer treatment, could be studied more comprehensively, thus contributing to the understanding of how extracellular conditions correlate with cancer cell proliferation and/or migration. The possibility to combine two or more Nbs with different biosensors might even broaden their application in multi-parametric imaging approaches, a technique highly desired in fluorescence microscopy (Carlson and Campbell, 2009). Finally, the biosensors proposed here can be further extended to monitor the effects of other factors such as cytokines, growth factors, hormones, ions, and metabolites that are known to determine how cells respond to their environment, mediating cell-to-cell and cell-to-matrix communication (Yang et al., 2019).

By flexibly linking two functionally independent modules, the herein presented principle of Nb-fused biosensors can be transferred to a range of signaling mediators or events utilizing appropriate FRET chromophores and Nbs that display suitable affinity and specificity for their target epitopes. In particular Nbs against a surface marker of solid tumor cells such as epidermal growth factor receptor (EGFR) (Roovers et al., 2007), prostate-specific membrane antigen (PSMA) (Evazalipour et al., 2014), carcinoembryonic antigen (CEA) (Cortez-Retamozo et al., 2004) or directly targeting components of the ECM such as the EIIIB domain of fibronectin (Jailkhani et al., 2019) may be promising candidates for the design of additional Nb-fused biosensors in the future.

In summary, the design and application of Nb-fused biosensors as demonstrated in this study address the growing need for reliable reagents to accurately determine analytes and their changes close to cell surfaces in the extracellular space. We assume that these new tools could pave the way to a better understanding of the cell-to-cell and cell-to-matrix communication elicited by ions, metabolites, and other signaling factors. Such versatile probes will open up new possibilities for the reliable investigation of extracellular analytes in advanced 3D cell models or even *in vivo* systems.

## MATERIAL & METHODS

### Cloning of Nb-fused biosensors

For the construction of the Nb-fused biosensors, the cDNA of SPOT-Nb (kindly provided by ChromoTek, Martinsried, Germany) and HER2-Nb 2Rs15d (Vaneycken *et al*., 2011) obtained as a gene synthesis product from Thermo Fisher (Schwerte, Germany) were used. The biosensor moieties used were the GEPII 1.0 potassium sensor (Bischof *et al*., 2017), the pH reporter pH-Lemon(Burgstaller *et al*., 2019), and the glucose sensor FLII12Pglu700µd6 (further referred to as FLII) (Takanaga *et al*., 2008). GEPII 1.0 consists of mseCFP, the potassium-binding domain (Kbp), and cpV, reporting K^+^ alterations by a conformational change of the Kbp (Bischof *et al*., 2017). Within pH-Lemon, the pH stable mTurquoise2 is directly connected to the pH-sensitive EYFP via a flexible linker and responds to pH changes due to the differential pH sensitivities of the mTurquoise2 (pKa 3.1) and EYFP (pKa 6.9) (Burgstaller *et al*., 2019; Goedhart et al., 2012; Ormö et al., 1996). Within FLII, the FRET donor ECFP is flanked by parts of the glucose-binding domain MglB on each end, while the FRET acceptor is located on the C-terminal end (Deuschle *et al*., 2005; Takanaga *et al*., 2008).

All expression constructs of Nb-fused biosensors were designed with the Nb at the N- and the sensor moiety at the C-terminus. Therefore the cDNA of the Nbs were genetically fused to the sensor thereby including a flexible (Gly_4_Ser)_4_ linker. Subsequently, SPOT-Nb- and HER2-Nb-containing constructs were cloned into a pET28a(+) vector (Merck-Millipore, Darmstadt, Germany), thereby adding a C-terminal hexahistidine tag (His_6_-tag) using conventional, PCR and restriction enzyme-based cloning. Finale expression constructs were subcloned in NEB 5- alpha competent *E.coli* cells (New England Biolabs, Ipswich, US) followed by Sanger sequencing (Microsynth, Göttingen, Germany).

### Protein expression

To achieve a high fraction of soluble biosensors SPOT-Nb-GEPII 1.0, SPOT-Nb-pH-Lemon, SPOT-Nb-FLII, HER2-Nb-GEPII 1.0, and HER2-Nb-pH-Lemon were transformed into chemically competent *E.coli* Arctic Express (DE3) (Agilent Technologies, Waldbronn, Germany)) following the manufactureŕs guidelines. Briefly, a 50 mL over-night culture containing 25 µg/mL kanamycin (Carl Roth GmbH, Karlsruhe, Germany) and G418 (Sigma Aldrich, Schnelldorf, Germany) was inoculated with one single colony. The over-night culture was transferred to 0,2 L main culture resulting in a starting OD_600_ of 0.2. Protein expression was induced at an OD_600_ of 0.8 using 0.5 mM IPTG (Carl Roth GmbH) followed by incubation for 24 h at 12°C. Subsequently, the cells were harvested (6000 x g, 10min, 4°C), resuspended in IMAC binding buffer (15 mM imidazole, 500 mM NaCl, pH 7.6 in PBS) containing 2 mM phenylmethylsulfonylfluorid (PMSF, AppliChem GmbH, Darmstadt, Germany), flash-frozen in liquid nitrogen and stored at -80°C. For the production of HER2-Nb, the expression construct was transformed into *E.coli* BL21 (DE3) following the manufactureŕs instructions. Briefly, one single colony was used to inoculate a 50 mL over-night culture containing 100 µg/mL ampicillin. The overnight culture was transferred to the 1 L main culture resulting in a starting OD_600_ of 0.2. Protein expression was induced at an OD_600_ of 0.8 using 1 mM IPTG followed by incubation for 20 h at RT. Cells were resuspended in IMAC binding buffer (15 mM imidazole, 500 mM NaCl, pH 7.6 in PBS) after harvesting (6000 x g, 10 min, 4°C) containing 2 mM PMSF, flash-frozen in liquid nitrogen and stored at -80°C.

### Protein purification

For protein purification, the bacterial cells were thawed and 0.16 mg/mL lysozyme (AppliChem GmbH, Darmstadt, Germany), 0.16 mg/mL DNAseI (ThermoFisher Scientific, Waltham, MA, USA) and protease inhibitor mix B (SERVA Electrophoresis GmbH, Heidelberg, Germany) were added, followed by 2 rounds (15 min each) of sonication, interrupted by 60 min incubation in a rotary shaker at 4°C. After harvesting (45 min, 18000 x g, 4°C), the supernatant of Nb-biosensors was collected, and the pellet was dissolved in 2 mL of 2 M imidazole to extract residual proteins. After sonication (4 x 30 sec) and harvesting (45 min, 18000 x g, 4°C), both supernatants were combined, filtered (0.45 µm) and diluted with 500 mM NaCl in PBS to a final imidazole concentration of 15 mM and a pH of 7.6. The solution was loaded onto a 5 mL HisTrap column (Cytiva, Massachusetts, US), washed with 20 column volumes of washing buffer (15 mM imidazole, 500 mM NaCl, pH 7.6 in PBS), followed by protein elution (500 mM imidazole, 500 mM NaCl, pH 7.4 in PBS) of individual fractions. Fractions showing high amounts of the target protein and low impurities (as determined using SDS-PAGE) were combined and concentrated using an Amicon concentrator tube (30 kDa MWCO for the Nb- fused biosensors, 3 kDa MWCO for HER2-Nb) (Merck-Millipore). A final volume < 5 mL was loaded onto a SEC column (HiLoad 200pg, 16&600 for Nb-fused biosensors; HiLoad 75pg, 26&600 for HER2-Nb) (Cytiva) and eluted with HEPES buffer (10 mM HEPES, 150 mM NaCl, pH 7.4 in ddH_2_0). SEC fractions containing full length proteins were concentrated to a concentration of ∼1 mg/mL as determined using NanoDrop (ThermoFisher). After the addition of glycerol to a final concentration of 2% (v/v), proteins were aliquoted, flash frozen and stored at -80°C until usage.

For quality control, all purified proteins were analyzed via SDS-PAGE according to standard procedures. Therefore, protein samples were denaturized (5 min, 95°C) in 2x SDS-sample buffer containing 60 mM Tris/HCl, pH 6.8; 2% (w/v) SDS; 5% (v/v) 2-mercaptoethanol, 10% (v/v) glycerol, 0.02% bromphenole blue. All proteins were visualized by InstantBlue Coomassie (Expedeon) staining. For immunoblotting, proteins were transferred to nitrocellulose membrane (GE Healthcare, Chicago, US) and detection was performed using Cy-5 labelled anti-VHH antibody (Cy™5 AffiniPure Goat Anti-Alpaca IgG, VHH domain, Jackson ImmunoResearch, UK) and a Typhoon Trio scanner (GE-Healthcare, excitation 633 nm, emission filter settings 670 nm BP 30).

### BioLayer Interferometry

To determine the binding affinity of purified SPOT-fused biosensors, biolayer interferometry (BLI) was performed on an Octet device (Sartorius GmbH, Germany) according to the manufactureŕs guidelines. Briefly, biotinylated SPOT peptide (Intavis, Tuebingen, Germany) loaded onto Streptavidin-coated sensor tips. With different concentrations of SPOT-Nb-GEPII 1.0 and SPOT-Nb-pH-Lemon (6.25, 12.5, 25 and 50 nM) and SPOT-Nb-FLII (12.5, 25, 50 and 100 nM) diluted in Octet buffer (0.1 % BSA (v/v), 0.02% Tween 20 (v/v) in PBS) four association/dissociation runs were performed to determine the association and dissociation rate constant.

### Cell culture, transfection, and assessment of cell viability

HEK293, MCF7 and SkBr3 cells were purchased from ATCC (Virginia, US). All cell lines were cultivated at 37°C and 5% CO_2_ in a humidified incubator using DMEM (ThermoFisher) + 10% FCS. For fluorescence widefield microcopy HEK293 cells were cultivated on 30 mm circular glass slides, while cells were seeded into 96 well imaging plates (Greiner, Kremsmünster, Austria) for confocal microscopy experiments. For immobilization of SPOT-Nb-sensors on HEK293 cells, the cells were transfected with a plasmid encoding a GPI-anchored SPOTtag (GPI-SPOT) using Lipofectamine 2000 reagent (ThermoFisher) according to the manufactureŕs instructions 2 days before experiments, followed by the removal of the transfection reagent after 6 h. For transient HER2 overexpression in HEK293 cells, the cells were transfected with a plasmid encoding human HER2 wildtype (Li et al., 2004)(Addgene plasmid #16257) the day before measurement with removal of the transfection reagent after 6 hours. Primary hippocampal neurons were isolated from early postnatal P0 C57BL/6N mice. Following preparation in dissection medium (HBSS with 1% sodium pyruvate, 1% HEPES (1 M) and 0.5% glucose (1 M)) hippocampal tissue was digested for 20 min in 2.5% trypsin at 37 °C. After repeated washing in dissection medium remaining tissue pieces were thoroughly triturated. Dissociated neurons were seeded at a density of 110,000 cells in ibidi 12 mm- Culture-Inserts (ibidi GmbH) on poly-L-Lysine coated 30 mm circular glass slides in 6 well plates. After cell attachment in plating medium (BME with 1% sodium pyruvate, 1% Glutamax, 10% FBS, 1% penicillin/streptomycin (P/S)) hippocampal neurons were cultured in maintenance medium (Neurobasal (NB) medium with 2% B27, 1% sodium pyruvate, 1% P/S) for 8 days. At DIV 5 20% (v/v) of the medium was exchanged with fresh NB medium. Neuronal cultures were transfected with GPI-SPOT using 0,32 µL lipofectamine and 1,6 µg DNA diluted in 160 µL OPTI-MEM (2x80 µL). 160µL culture medium (from 0,5 mL total volume) was removed and 160 µL of the transfection mix was added. On two consecutive days (24 and 48 h after the start of transfection), 3/4 of the medium was replaced to dilute the transection mix. Cells were analyzed 5 days after transfection.

Cell viability over time of control cells (no incubation with Nb) and after immobilization of HER2- Nb and HER2-Nb-GEPII 1.0 was assessed using 3-(4,5-Dimethylthiazol-2-yl)-2,5-Diphenyltetrazolium Bromide (MTT) reagent (ThermoFisher). After immobilization of HER2-Nb and HER2-Nb-GEPII 1.0 on the cell surface, cells were cultivated for either 3, 6, 24 or 48 h in DMEM + 10% FCS prior to the assay. MTT assay was performed according to manufacturer’s instructions. Absorbance at 540 nm was recorded using a Tecan Infinite 200 PRO plate reader. For all time-points, cells without immobilized HER2-Nb or HER2-Nb-biosensor served as a 100% viability control.

### Immobilization of Nb-biosensors

For immobilization of SPOT-Nb-GEPII 1.0, SPOT-Nb-pH-Lemon, SPOT-Nb-FLII, HER2-Nb- GEPII 1.0 and HER2-Nb-pH-Lemon, the proteins were diluted to a final concentration of 2 µM in a physiological buffer (138 mM NaCl, 5 mM KCl, 2 mM CaCl_2_, 1 mM MgCl_2_, 10 mM D-glucose, 10 mM HEPES, pH 7.4 in ddH_2_0) to a final volume of 1 mL (for 6 –well format) or 0.2 mL (for 96-well format). Cells were washed once with PBS and incubated with the Nb-based biosensor solution for 20 - 30 min at RT in the dark. After immobilization, the cells were washed 3 times with physiological buffer and immediately used for measurements.

### Fluorescence microscopy

For high-resolution fluorescence microscopy an ImageXpress Micro Confocal Microscope (Molecular Devices, California, US) equipped with a 40X water immersion objective was used. Cells were imaged in a physiological buffer, acquiring DAPI, mseCFP/mTurquoise2 and cpV/EYFP images. For FRET-based live-cell imaging experiments, a Zeiss Axio Observer Z1 equipped with a 40x oil immersion objective (EC “Plan-Neofluar” 40x/1,30 Oil M27) (Zeiss, Oberkochen, Germany) and connected to an OptoSplit II emission image splitter (Cairn Research, Faversham, UK) equipped with a T505lpxr (AHF Analysentechnik, Tübingen, Germany) was used. For illumination, an LedHUB LED light-engine equipped with a 455 nm LED and a 505 - 600 nM LED (Omicron, Dudenhofen, Germany) with 427/10 nm and 510/10 nm excitation filters (AHF Analysentechnik) was employed. Dichroic and emission filter in the microscope were as follows: 459/526/596 and 475/543/702 (AHF Analysentechnik). Image acquisition and microscope control were performed using VisiView software (Visitron Systems GmbH, Puchheim, Germany). For buffer exchange, a gravity-based perfusion system (NGFI GmbH, Graz, Austria) was connected to a PC30 perfusion chamber (NGFI GmbH).

### Functional characterization of Nb-fused biosensors

For the titration of different K^+^, pH and glucose concentrations, the physiological buffer was modified accordingly. Briefly, buffers for K^+^ titrations consisted of 2 mM CaCl_2_, 1 mM MgCl_2_, 10 mM HEPES, 10 mM glucose, pH 7.4 with different concentration of KCl to obtain 0, 0.3, 1, 3, 10, 30, and 100 mM K^+^. To maintain buffer osmolarity, the concentration of NaCl was adjusted. Buffers for pH titrations consisted of physiological buffers pH values were adjusted using N-Methyl-D-glucamin (NMDG) or HCl to obtain pH 4, 5, 6, 7, 7.5, 8 and 9. Buffers for glucose titrations consisted of physiological buffer. Different glucose concentrations were used to obtain 0, 0.1, 0.3, 1, 3, 10, 30 and 100 mM glucose.

### Visualization of neuronal K^+^ release

For visualization of K^+^ release, hippocampal neurons growing on 30 mm circular glass slides were inserted into the PC30 perfusion chamber and connected to the perfusion system (NGFI GmbH). Experiments were performed using the Zeiss Axio Observer Z1 microscope. 2 µM SPOT-Nb-GEPII 1.0 was immobilized on the cell surface for 30 min in physiological buffer. To remove unbound sensor protein after immobilization, the cells were shortly perfused with K^+^- free buffer. Subsequently, the perfusion was stopped, and 50 µL of the pure buffer without glutamate was added by pipetting, followed by a glutamate bolus (50 µL, 100 µM final concentration). Following the K^+^ increase upon glutamate addition, the perfusion was started to remove extracellular K^+^ using K^+^-free physiological buffer, followed by perfusion with a 100 mM K^+^ buffer to determine the maximal sensor response.

### Data representation and statistical analysis

Data analysis was performed using Excel software (Microsoft, New Mexico, US). Statistical analysis was performed using GraphPad Prism 5 software (GraphPad Software, San Diego, US). Data were analyzed for normal distribution using D’Agostino & Pearson omnibus normality test. P values < 0.05 were considered significant. Statistical tests used for the analysis of respective panels are indicated in the figure legend.

## Data availability

The data that support the findings of this study are available from the corresponding author upon reasonable request.

## Author contributions

S.B., R.L. and U.R. conceived the study and the research design. S.B., T.R.W., H.B., S.Bu, D.S. P.D.K. D.Sk. performed protein expression, biochemical characterization and cellular experiments including imaging studies and FRET measurements. S.B., H.B. T.R.W. and D.S. analyzed the data. A.P., H.B., R.M. contributed new reagents or analytic tools. R.L. and U.R. supervised the study. S.B., R.L. and U.R. drafted the manuscript. All authors critically read the manuscript.

## Acknowledgments

Work in the laboratory of RL was supported by DFG research grants LU 1490/8-1 and LU 1490/10-1. RL acknowledges financial support from the Doktor Robert Pfleger-Stiftung, the ICEPHA Graduate Program ‘‘Membrane-associated Drug Targets in Personalized Cancer Medicine’’, and Cyclerion Therapeutics Inc. HB is a fellow of the FWF funded Erwin-Schrödinger-Program, project number J4457. For the purpose of open access, the author has applied a CC BY public copyright license to any Author Accepted Manuscript version arising from this submission. Work in the laboratory of UR was supported by the State Ministry of Baden-Württemberg for Economic Affairs, Labour and Housing Construction (FKZ 3-4332.62-NMI/68).

## Supplementary Figures

**Supplementary Figure 1.**
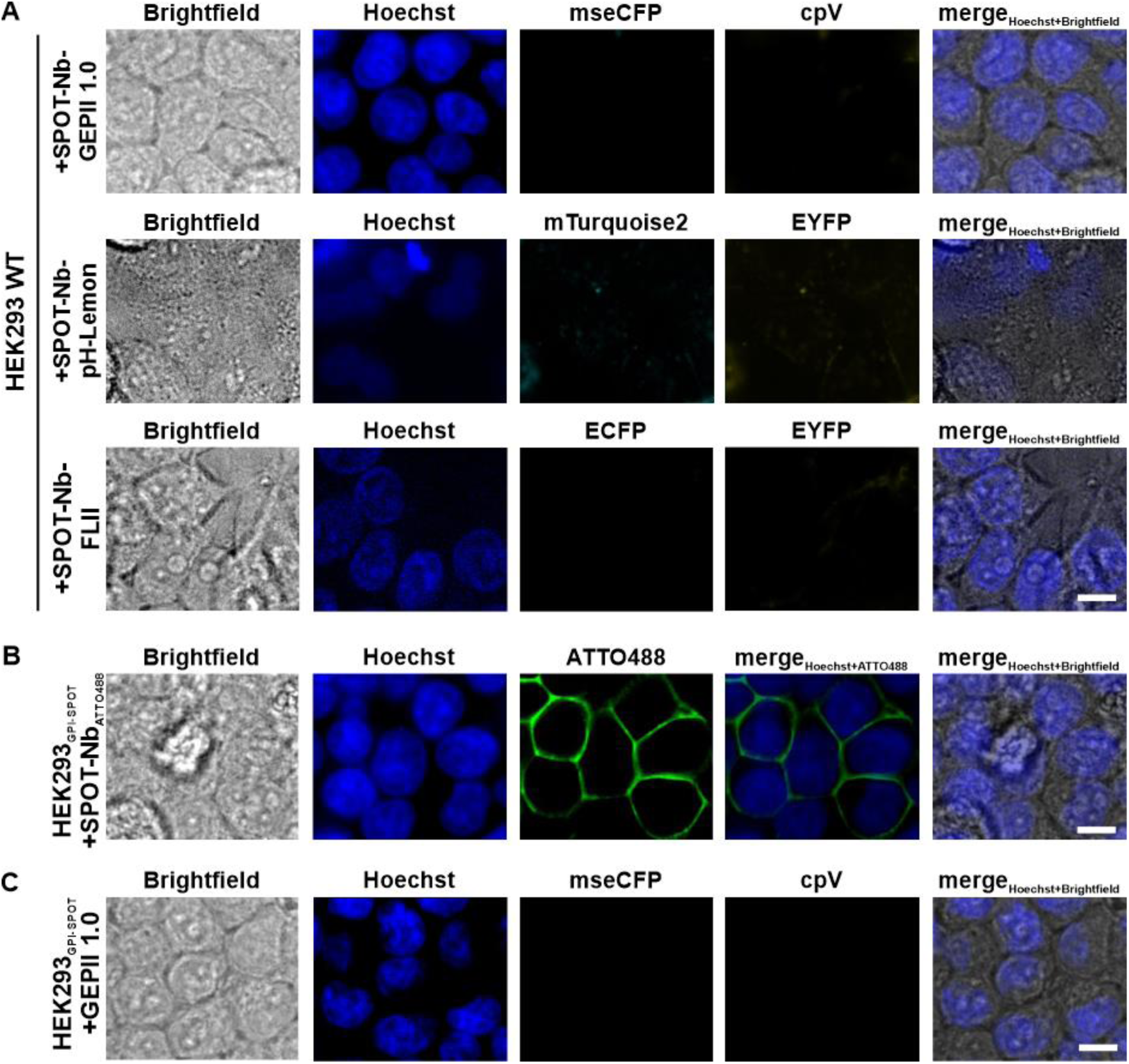
SPOT-Nb-biosensors do not bind unspecifically to the plasma membrane. Representative confocal images of (**A**) live HEK293 wildtype (WT) cells following incubation with SPOT-Nb-GEPII 1.0 (upper panel), SPOT-Nb-pH Lemon (middle panel) and SPOT-Nb-FLII (lower panel. (**B**) Live imaging of HEK293 cells expressing GPI-anchored SPOT-tag (GPI-SPOT) on the plasma membrane after staining with the fluorescently labeled SPOT-Nb (SPOT-Nb_ATTO488_) and (**C**) of live HEK293 GPI-SPOT cells upon incubation with GEPII 1.0 biosensor lacking the SPOT-Nb. (**A – C**) Scale bar 10 µm, n= 4 experiments for (**A**, upper & middle row) and (**B**); n= 2 experiments for (**A**, lower row) and (**C**).

**Supplementary Figure 2.**
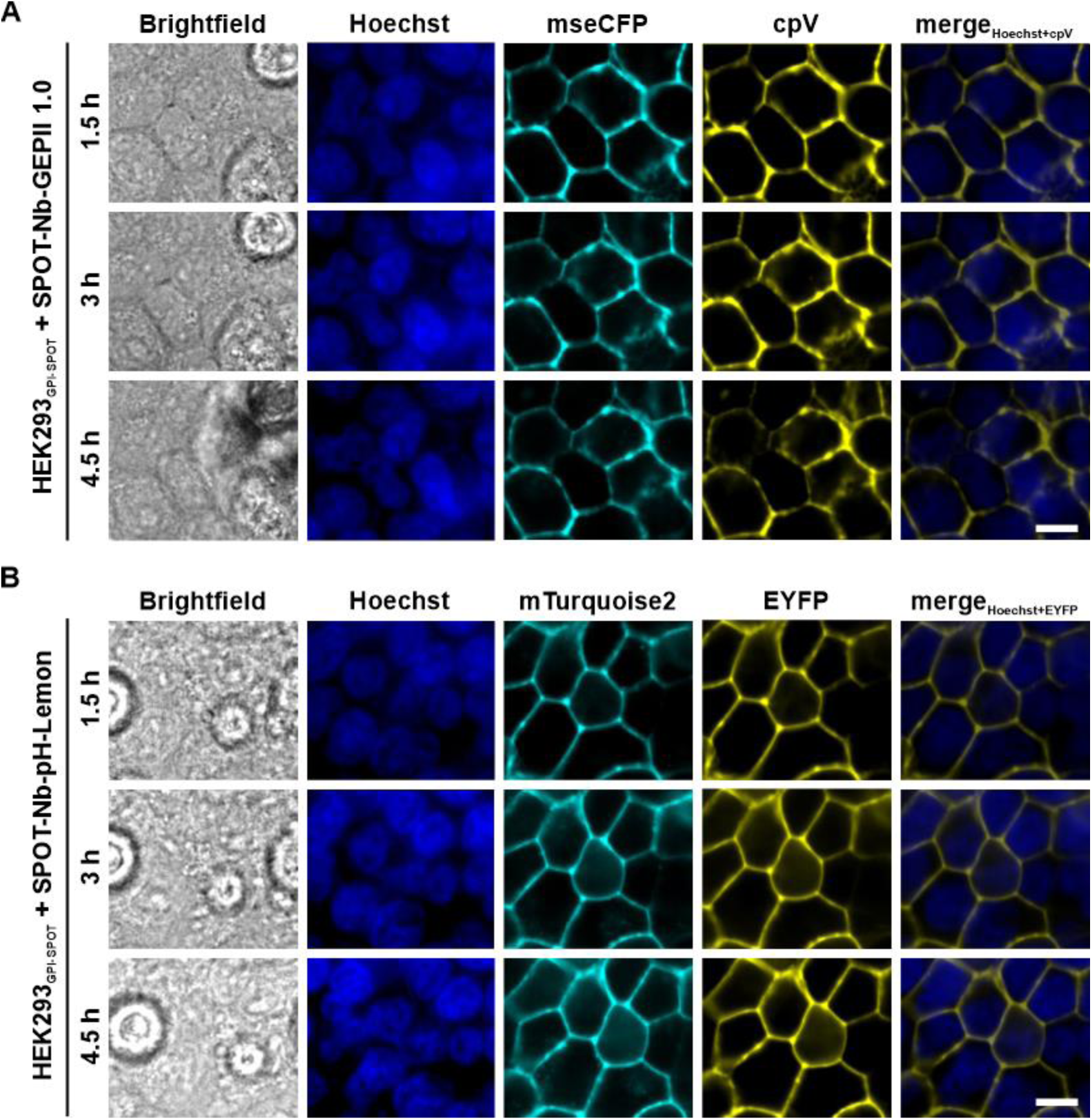
Long term binding of SPOT-Nb-biosensors to the plasma membrane of living cells. Time-lapse confocal microscopy of HEK293 GPI-SPOT expressing cells following incubation with SPOT-Nb-GEPII 1.0 (**A**) or SPOT-Nb-pH-Lemon (**B**). Representative images of n=4 experiments at indicated time points are shown. Scale bar 10 µm.

**Supplementary Figure 3.**
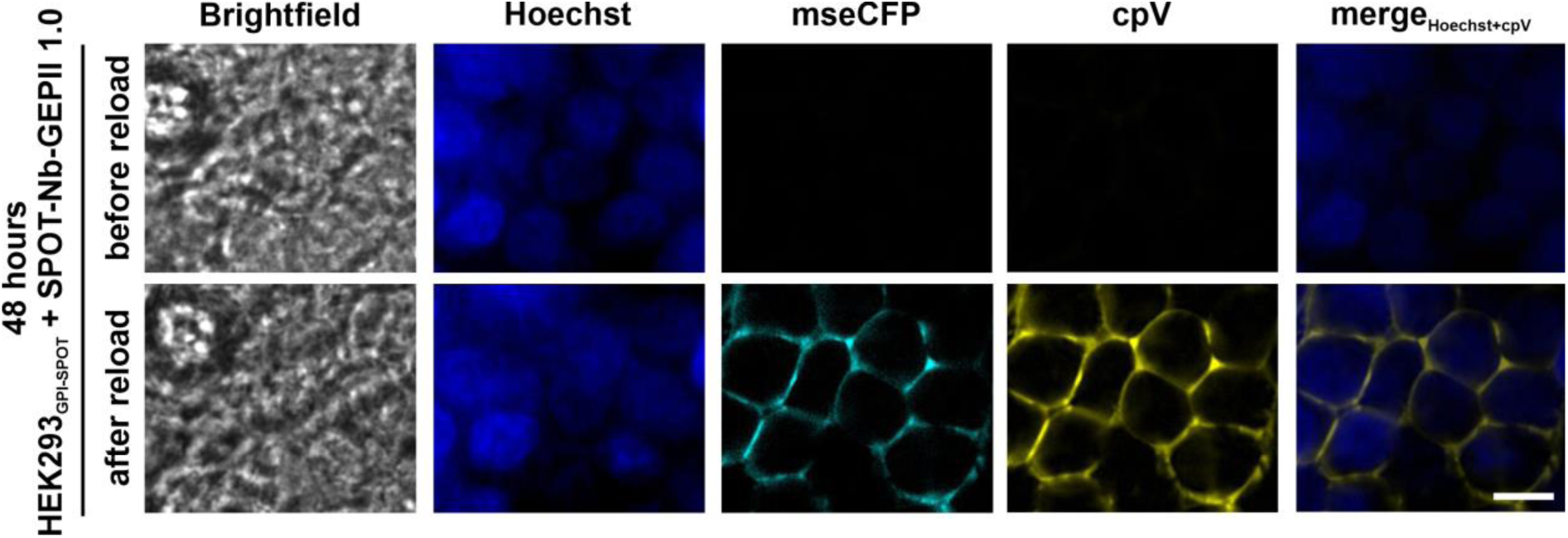
Re-loading of the GPI-SPOT anchor with SPOT-Nb-GEPII 1.0 at the plasma membrane. Representative confocal images of HEK293 GPI-SPOT cells 48 h after incubation with SPOT-Nb-GEPII 1.0 (upper row) or upon re-loading of SPOT-Nb-GEPII 1.0 (lower row). Scale bar 10 µm.

**Supplementary Figure 4.**
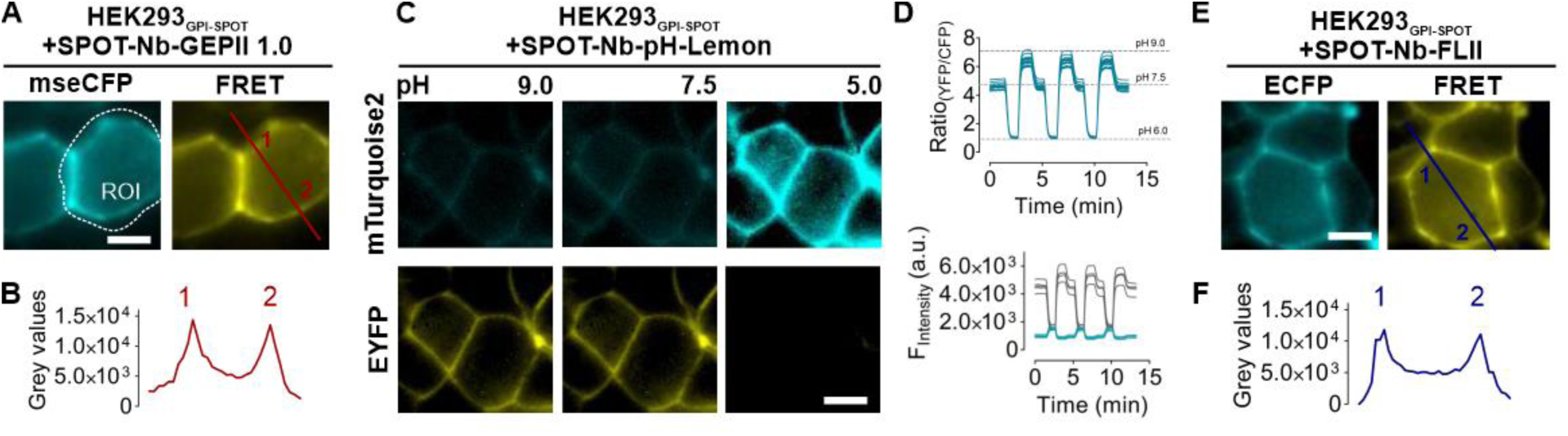
Widefield FRET-imaging of SPOT-Nb-biosensors. (**A**) Representative widefield images of HEK293 GPI-SPOT cells displaying the mseCFP (left panel) and the FRET signal (right panel) derived from SPOT-Nb-GEPII 1.0 binding. An exemplary region (ROI, white dotted line) used for measuring the FRET response is indicated. (**B**) Intensity profile of a line scan (red) crossing the plasma membrane. Peaks (1 and 2) in (**B**) correspond with the respective labelling of FRET image in (**A**). (**C**) Representative widefield images of HEK293 GPI-SPOT cells displaying either mTurquoise2 (upper row) or EYFP signal (lower row) derived from bound SPOT-Nb-pH-Lemon in response to alkaline pH (9.0), neutral pH (7.5) and acidic pH (5.0). (**D**) Ratio signals of individual cells in response to repetitive pH alterations (switching between pH 7.5, 6.0, and 9.0, upper panel). Respective single wavelength traces (EYFP, grey and mTurquoise2, blue) upon pH alterations (lower panel). (**E**) Representative widefield images of HEK293 GPI-SPOT cells displaying ECFP (left panel) or FRET signal (right panel) derived from bound SPOT-Nb-FLII. (**F**) Intensity profile of a line scan (blue) crossing the plasma membrane. Peaks (1 and 2) in (**F**) correspond with the respective labelling of FRET image in (**E**). (**A,C,E**) Scale bar 10 µM, (**A,C,D,E**) n= 3 experiments from biological replicates.

**Supplementary Figure 5.**
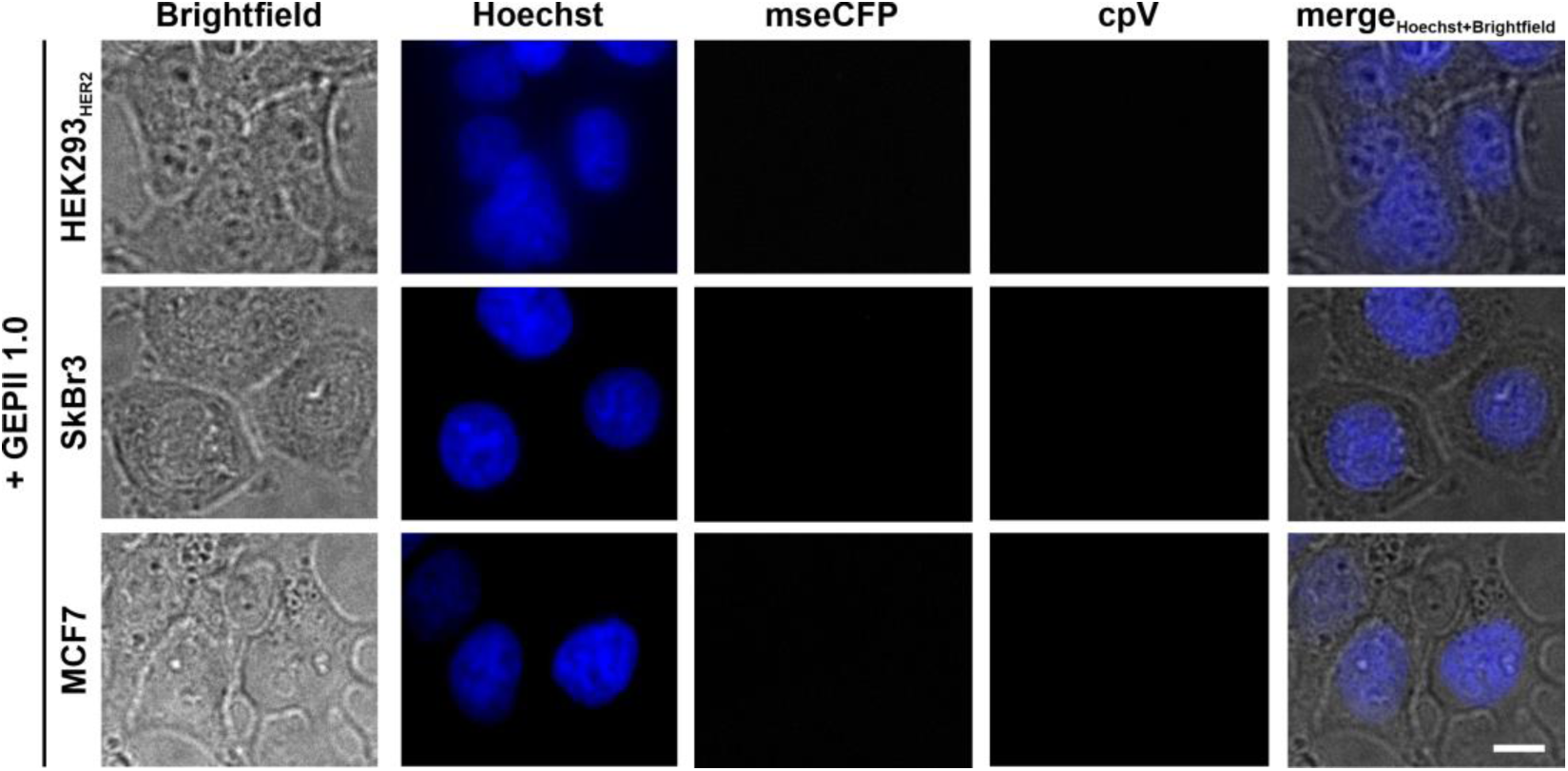
Purified GEPII 1.0 lacking the HER2-Nb does not bind to the cell surface of cancer cells. Representative confocal live imaging of HEK293 cells transiently overexpressing HER2 (HEK293_HER2_, upper row), SkBr3 cells (middle row) and MCF7 cells (lower row) after addition of purified GEPII 1.0 lacking the HER2-Nb are shown. Scale bar 10 µm, n= 2 experiments.

**Supplementary Figure 6.**
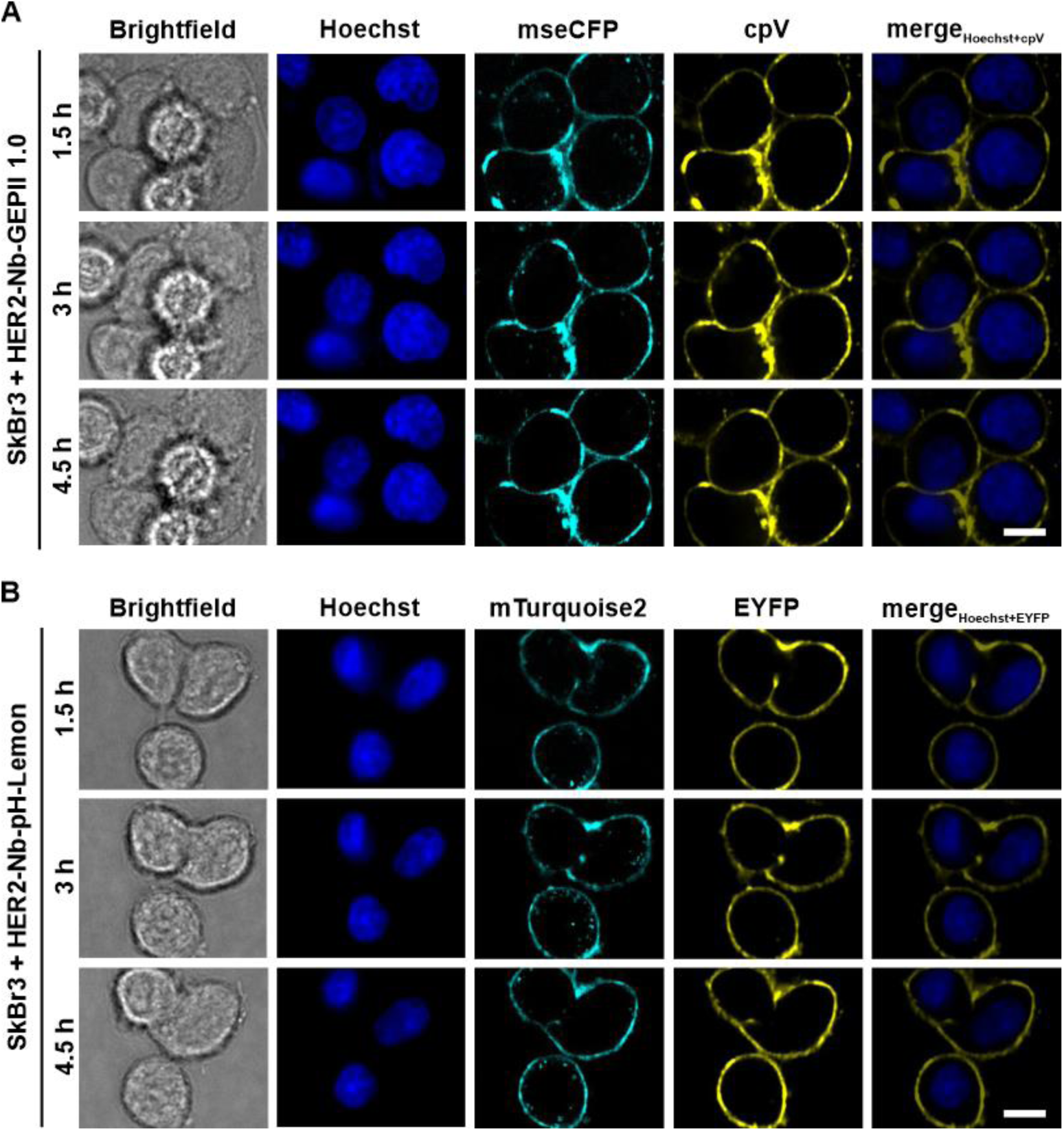
Long term binding of HER2-Nb-biosensors to the plasma membrane of living cancer cells. Time-lapse microscopy of SkBr3 cells following incubation with HER2-Nb-GEPII 1.0 (**A**) or HER2-Nb-pH-Lemon (**B**). Representative images at indicated time points are shown. Scale bar 10 µm, n= 2 experiments

**Supplementary Table 1:** Binding affinities of SPOT-Nb-biosensors determined using biolayer interferometry.

| Nbs | $K_D$ [nM] | $k_{on}$ [$M^{-1}s^{-1}$] | $k_{off}$ [$s^{-1}$] |
| --- | --- | --- | --- |
| SPOT-Nb-GEPII 1.0 | $0.73 \pm 0.09$ | $3.60 \pm 0.01 \times 10^4$ | $2.62 \pm 0.003 \times 10^{-4}$ |
| SPOT-Nb-pH-Lemon | $14.3 \pm 0.06$ | $2.21 \pm 0.008 \times 10^4$ | $3.16 \pm 0.003 \times 10^{-4}$ |
| SPOT-Nb-FLII | $73.4 \pm 1.05$ | $3.56 \pm 0.05 \times 10^3$ | $2.65 \pm 0.003 \times 10^{-4}$ |

## REFERENCES

Anderson, N.M., and Simon, M.C. (2020). The tumor microenvironment. Curr Biol 30, R921–r925. 10.1016/j.cub.2020.06.081.

Bischof, H., Burgstaller, S., Waldeck-Weiermair, M., Rauter, T., Schinagl, M., Ramadani-Muja, J., Graier, W.F., and Malli, R. (2019). Live-Cell Imaging of Physiologically Relevant Metal Ions Using Genetically Encoded FRET-Based Probes. Cells 8. 10.3390/cells8050492.

Bischof, H., Rehberg, M., Stryeck, S., Artinger, K., Eroglu, E., Waldeck-Weiermair, M., Gottschalk, B., Rost, R., Deak, A.T., Niedrist, T., et al. (2017). Novel genetically encoded fluorescent probes enable real-time detection of potassium in vitro and in vivo. Nature Communications 8, 1422. 10.1038/s41467-017-01615-z.

Braun, M.B., Traenkle, B., Koch, P.A., Emele, F., Weiss, F., Poetz, O., Stehle, T., and Rothbauer, U. (2016). Peptides in headlock--a novel high-affinity and versatile peptide-binding nanobody for proteomics and microscopy. Sci Rep 6, 19211. 10.1038/srep19211.

Burgstaller, S., Bischof, H., Gensch, T., Stryeck, S., Gottschalk, B., Ramadani-Muja, J., Eroglu, E., Rost, R., Balfanz, S., Baumann, A., et al. (2019). pH-Lemon, a Fluorescent Protein-Based pH Reporter for Acidic Compartments. ACS Sens 4, 883–891. 10.1021/acssensors.8b01599.

Burgstaller, S., Bischof, H., Matt, L., and Lukowski, R. (2022). Assessing K(+) ions and K(+) channel functions in cancer cell metabolism using fluorescent biosensors. Free Radic Biol Med 181, 43–51. 10.1016/j.freeradbiomed.2022.01.026.

Burgstaller, S., Bischof, H., Rauter, T., Schmidt, T., Schindl, R., Patz, S., Groschup, B., Filser, S., van den Boom, L., Sasse, P., et al. (2021). Immobilization of Recombinant Fluorescent Biosensors Permits Imaging of Extracellular Ion Signals. ACS Sens 6, 3994–4000. 10.1021/acssensors.1c01369.

Busco, G., Cardone, R.A., Greco, M.R., Bellizzi, A., Colella, M., Antelmi, E., Mancini, M.T., Dell’Aquila, M.E., Casavola, V., Paradiso, A., and Reshkin, S.J. (2010). NHE1 promotes invadopodial ECM proteolysis through acidification of the peri-invadopodial space. Faseb j 24, 3903–3915. 10.1096/fj.09-149518.

Cao, X., Fang, L., Gibbs, S., Huang, Y., Dai, Z., Wen, P., Zheng, X., Sadee, W., and Sun, D. (2007). Glucose uptake inhibitor sensitizes cancer cells to daunorubicin and overcomes drug resistance in hypoxia. Cancer Chemother Pharmacol 59, 495–505. 10.1007/s00280-006-0291-9.

Carlson, H.J., and Campbell, R.E. (2009). Genetically encoded FRET-based biosensors for multiparameter fluorescence imaging. Curr Opin Biotechnol 20, 19–27. 10.1016/j.copbio.2009.01.003.

Comes, N., Serrano-Albarrás, A., Capera, J., Serrano-Novillo, C., Condom, E., Ramón, Y.C.S., Ferreres, J.C., and Felipe, A. (2015). Involvement of potassium channels in the progression of cancer to a more malignant phenotype. Biochim Biophys Acta 1848, 2477–2492. 10.1016/j.bbamem.2014.12.008.

Cortez-Retamozo, V., Backmann, N., Senter, P.D., Wernery, U., De Baetselier, P., Muyldermans, S., and Revets, H. (2004). Efficient cancer therapy with a nanobody-based conjugate. Cancer Res 64, 2853–2857. 10.1158/0008-5472.can-03-3935.

Costanza, B., Rademaker, G., Tiamiou, A., De Tullio, P., Leenders, J., Blomme, A., Bellier, J., Bianchi, E., Turtoi, A., Delvenne, P., et al. (2019). Transforming growth factor beta-induced, an extracellular matrix interacting protein, enhances glycolysis and promotes pancreatic cancer cell migration. Int J Cancer 145, 1570–1584. 10.1002/ijc.32247.

Depaoli, M.R., Bischof, H., Eroglu, E., Burgstaller, S., Ramadani-Muja, J., Rauter, T., Schinagl, M., Waldeck-Weiermair, M., Hay, J.C., Graier, W.F., and Malli, R. (2019). Live cell imaging of signaling and metabolic activities. Pharmacol Ther 202, 98–119. 10.1016/j.pharmthera.2019.06.003.

Deuschle, K., Okumoto, S., Fehr, M., Looger, L.L., Kozhukh, L., and Frommer, W.B. (2005). Construction and optimization of a family of genetically encoded metabolite sensors by semirational protein engineering. Protein Sci 14, 2304–2314. 10.1110/ps.051508105.

Ehinger, R., Kuret, A., Matt, L., Frank, N., Wild, K., Kabagema-Bilan, C., Bischof, H., Malli, R., Ruth, P., Bausch, A.E., and Lukowski, R. (2021). Slack K(+) channels attenuate NMDA-induced excitotoxic brain damage and neuronal cell death. Faseb j 35, e21568. 10.1096/fj.202002308RR.

Eil, R., Vodnala, S.K., Clever, D., Klebanoff, C.A., Sukumar, M., Pan, J.H., Palmer, D.C., Gros, A., Yamamoto, T.N., Patel, S.J., et al. (2016). Ionic immune suppression within the tumour microenvironment limits T cell effector function. Nature 537, 539–543. 10.1038/nature19364.

Evazalipour, M., D’Huyvetter, M., Tehrani, B.S., Abolhassani, M., Omidfar, K., Abdoli, S., Arezumand, R., Morovvati, H., Lahoutte, T., Muyldermans, S., and Devoogdt, N. (2014). Generation and characterization of nanobodies targeting PSMA for molecular imaging of prostate cancer. Contrast Media Mol Imaging 9, 211–220. 10.1002/cmmi.1558.

Feilmeier, B.J., Iseminger, G., Schroeder, D., Webber, H., and Phillips, G.J. (2000). Green fluorescent protein functions as a reporter for protein localization in Escherichia coli. J Bacteriol 182, 4068–4076. 10.1128/jb.182.14.4068-4076.2000.

Fiszer-Kierzkowska, A., Vydra, N., Wysocka-Wycisk, A., Kronekova, Z., Jarząb, M., Lisowska, K.M., and Krawczyk, Z. (2011). Liposome-based DNA carriers may induce cellular stress response and change gene expression pattern in transfected cells. BMC Mol Biol 12, 27. 10.1186/1471-2199-12-27.

Frantz, C., Karydis, A., Nalbant, P., Hahn, K.M., and Barber, D.L. (2007). Positive feedback between Cdc42 activity and H+ efflux by the Na-H exchanger NHE1 for polarity of migrating cells. J Cell Biol 179, 403–410. 10.1083/jcb.200704169.

Gallagher, F.A., Kettunen, M.I., Day, S.E., Hu, D.E., Ardenkjaer-Larsen, J.H., Zandt, R., Jensen, P.R., Karlsson, M., Golman, K., Lerche, M.H., and Brindle, K.M. (2008). Magnetic resonance imaging of pH in vivo using hyperpolarized 13C-labelled bicarbonate. Nature 453, 940–943. 10.1038/nature07017.

Goedhart, J., von Stetten, D., Noirclerc-Savoye, M., Lelimousin, M., Joosen, L., Hink, M.A., van Weeren, L., Gadella, T.W., Jr., and Royant, A. (2012). Structure-guided evolution of cyan fluorescent proteins towards a quantum yield of 93%. Nat Commun 3, 751. 10.1038/ncomms1738.

Hamers-Casterman, C., Atarhouch, T., Muyldermans, S., Robinson, G., Hamers, C., Songa, E.B., Bendahman, N., and Hamers, R. (1993). Naturally occurring antibodies devoid of light chains. Nature 363, 446–448. 10.1038/363446a0.

Han, J., Zhang, L., Guo, H., Wysham, W.Z., Roque, D.R., Willson, A.K., Sheng, X., Zhou, C., and Bae-Jump, V.L. (2015). Glucose promotes cell proliferation, glucose uptake and invasion in endometrial cancer cells via AMPK/mTOR/S6 and MAPK signaling. Gynecol Oncol 138, 668–675. 10.1016/j.ygyno.2015.06.036.

Hösli, L., Hösli, E., Landolt, H., and Zehntner, C. (1981). Efflux of potassium from neurones excited by glutamate and aspartate causes a depolarization of cultured glial cells. Neurosci Lett 21, 83–86. 10.1016/0304-3940(81)90062-8.

Huang, S., Tang, Y., Peng, X., Cai, X., Wa, Q., Ren, D., Li, Q., Luo, J., Li, L., Zou, X., and Huang, S. (2016). Acidic extracellular pH promotes prostate cancer bone metastasis by enhancing PC-3 stem cell characteristics, cell invasiveness and VEGF-induced vasculogenesis of BM-EPCs. Oncol Rep 36, 2025–2032. 10.3892/or.2016.4997.

Huber, S.M. (2013). Oncochannels. Cell Calcium 53, 241–255. 10.1016/j.ceca.2013.01.001.

Huntington, K.E., Louie, A., Zhou, L., Seyhan, A.A., Maxwell, A.W., and El-Deiry, W.S. (2022). Colorectal cancer extracellular acidosis decreases immune cell killing and is partially ameliorated by pH-modulating agents that modify tumor cell cytokine profiles. Am J Cancer Res 12, 138–151.

Jacobsen, L., Calvin, S., and Lobenhofer, E. (2009). Transcriptional effects of transfection: the potential for misinterpretation of gene expression data generated from transiently transfected cells. Biotechniques 47, 617–624. 10.2144/000113132.

Jähde, E., Glüsenkamp, K.H., and Rajewsky, M.F. (1990). Protection of cultured malignant cells from mitoxantrone cytotoxicity by low extracellular pH: a possible mechanism for chemoresistance in vivo. Eur J Cancer 26, 101–106. 10.1016/0277-5379(90)90290-a.

Jailkhani, N., Ingram, J.R., Rashidian, M., Rickelt, S., Tian, C., Mak, H., Jiang, Z., Ploegh, H.L., and Hynes, R.O. (2019). Noninvasive imaging of tumor progression, metastasis, and fibrosis using a nanobody targeting the extracellular matrix. Proceedings of the National Academy of Sciences 116, 14181–14190. doi:10.1073/pnas.1817442116.

Jain, R.K., Joyce, P.B., Molinete, M., Halban, P.A., and Gorr, S.U. (2001). Oligomerization of green fluorescent protein in the secretory pathway of endocrine cells. Biochem J 360, 645–649. 10.1042/0264-6021:3600645.

Kunz, P., Zinner, K., Mücke, N., Bartoschik, T., Muyldermans, S., and Hoheisel, J.D. (2018). The structural basis of nanobody unfolding reversibility and thermoresistance. Sci Rep 8, 7934. 10.1038/s41598-018-26338-z.

Li, Y.M., Pan, Y., Wei, Y., Cheng, X., Zhou, B.P., Tan, M., Zhou, X., Xia, W., Hortobagyi, G.N., Yu, D., and Hung, M.C. (2004). Upregulation of CXCR4 is essential for HER2-mediated tumor metastasis. Cancer Cell 6, 459–469. 10.1016/j.ccr.2004.09.027.

Liu, M., Li, L., Jin, D., and Liu, Y. (2021). Nanobody-A versatile tool for cancer diagnosis and therapeutics. Wiley Interdiscip Rev Nanomed Nanobiotechnol 13, e1697. 10.1002/wnan.1697.

Mello de Queiroz, F., Sánchez, A., Agarwal, J.R., Stühmer, W., and Pardo, L.A. (2012). Nucleofection induces non-specific changes in the metabolic activity of transfected cells. Mol Biol Rep 39, 2187–2194. 10.1007/s11033-011-0967-z.

Mohr, C.J., Gross, D., Sezgin, E.C., Steudel, F.A., Ruth, P., Huber, S.M., and Lukowski, R. (2019a). K(Ca)3.1 Channels Confer Radioresistance to Breast Cancer Cells. Cancers (Basel) 11. 10.3390/cancers11091285.

Mohr, C.J., Gross, D., Sezgin, E.C., Steudel, F.A., Ruth, P., Huber, S.M., and Lukowski, R. (2019b). KCa3.1 Channels Confer Radioresistance to Breast Cancer Cells. Cancers 11, 1285.

Mohr, C.J., Schroth, W., Mürdter, T.E., Gross, D., Maier, S., Stegen, B., Dragoi, A., Steudel, F.A., Stehling, S., Hoppe, R., et al. (2020). Subunits of BK channels promote breast cancer development and modulate responses to endocrine treatment in preclinical models. Br J Pharmacol. 10.1111/bph.15147.

Mohr, C.J., Steudel, F.A., Gross, D., Ruth, P., Lo, W.-Y., Hoppe, R., Schroth, W., Brauch, H., Huber, S.M., and Lukowski, R. (2019c). Cancer-Associated Intermediate Conductance Ca2+- Activated K+ Channel KCa3.1. Cancers 11, 109.

Muyldermans, S. (2013). Nanobodies: natural single-domain antibodies. Annu Rev Biochem 82, 775–797. 10.1146/annurev-biochem-063011-092449.

Muyldermans, S. (2021). Applications of Nanobodies. Annual Review of Animal Biosciences 9, 401–421. 10.1146/annurev-animal-021419-083831.

Namiki, S., Sakamoto, H., Iinuma, S., Iino, M., and Hirose, K. (2007). Optical glutamate sensor for spatiotemporal analysis of synaptic transmission. Eur J Neurosci 25, 2249–2259. 10.1111/j.1460-9568.2007.05511.x.

Olichon, A., and Surrey, T. (2007). Selection of genetically encoded fluorescent single domain antibodies engineered for efficient expression in Escherichia coli. J Biol Chem 282, 36314–36320. 10.1074/jbc.M704908200.

Ormö, M., Cubitt, A.B., Kallio, K., Gross, L.A., Tsien, R.Y., and Remington, S.J. (1996). Crystal Structure of the *Aequorea victoria* Green Fluorescent Protein. Science 273, 1392–1395. doi:10.1126/science.273.5280.1392.

Pardo, L.A., and Stühmer, W. (2014). The roles of K+ channels in cancer. Nature Reviews Cancer 14, 39–48. 10.1038/nrc3635.

Pedersen, A.K., Mendes Lopes de Melo, J., Mørup, N., Tritsaris, K., and Pedersen, S.F. (2017). Tumor microenvironment conditions alter Akt and Na+/H+ exchanger NHE1 expression in endothelial cells more than hypoxia alone: implications for endothelial cell function in cancer. BMC Cancer 17, 542. 10.1186/s12885-017-3532-x.

Peruzzo, R., and Szabo, I. (2019). Contribution of Mitochondrial Ion Channels to Chemo-Resistance in Cancer Cells. Cancers (Basel) 11. 10.3390/cancers11060761.

Prole, D.L., and Taylor, C.W. (2019). A genetically encoded toolkit of functionalized nanobodies against fluorescent proteins for visualizing and manipulating intracellular signalling. BMC Biology 17, 41. 10.1186/s12915-019-0662-4.

Roovers, R.C., Laeremans, T., Huang, L., De Taeye, S., Verkleij, A.J., Revets, H., de Haard, H.J., and van Bergen en Henegouwen, P.M.P. (2007). Efficient inhibition of EGFR signalling and of tumour growth by antagonistic anti-EGFR Nanobodies. Cancer Immunology, Immunotherapy 56, 303–317. 10.1007/s00262-006-0180-4.

Sankaranarayanan, S., De Angelis, D., Rothman, J.E., and Ryan, T.A. (2000). The use of pHluorins for optical measurements of presynaptic activity. Biophys J 79, 2199–2208. 10.1016/s0006-3495(00)76468-x.

Shen, Y., Rosendale, M., Campbell, R.E., and Perrais, D. (2014). pHuji, a pH-sensitive red fluorescent protein for imaging of exo-and endocytosis. J Cell Biol 207, 419–432. 10.1083/jcb.201404107.

Som, A., Bloch, S., Ippolito, J.E., and Achilefu, S. (2016). Acidic extracellular pH of tumors induces octamer-binding transcription factor 4 expression in murine fibroblasts in vitro and in vivo. Scientific Reports 6, 27803. 10.1038/srep27803.

Soroceanu, L., Manning, T.J., and Sontheimer, H. (1999). Modulation of Glioma Cell Migration and Invasion Using Cl^−^ and K^+^ Ion Channel Blockers. The Journal of Neuroscience 19, 5942–5954. 10.1523/jneurosci.19-14-05942.1999.

Steudel, F.A., Mohr, C.J., Stegen, B., Nguyen, H.Y., Barnert, A., Steinle, M., Beer-Hammer, S., Koch, P., Lo, W.Y., Schroth, W., et al. (2017). SK4 channels modulate Ca(2+) signalling and cell cycle progression in murine breast cancer. Mol Oncol 11, 1172–1188. 10.1002/1878-0261.12087.

Takanaga, H., Chaudhuri, B., and Frommer, W.B. (2008). GLUT1 and GLUT9 as major contributors to glucose influx in HepG2 cells identified by a high sensitivity intramolecular FRET glucose sensor. Biochim Biophys Acta 1778, 1091–1099. 10.1016/j.bbamem.2007.11.015.

Tantama, M., Martínez-François, J.R., Mongeon, R., and Yellen, G. (2013). Imaging energy status in live cells with a fluorescent biosensor of the intracellular ATP-to-ADP ratio. Nature Communications 4, 2550. 10.1038/ncomms3550.

Thews, O., Nowak, M., Sauvant, C., and Gekle, M. (2011). Hypoxia-Induced Extracellular Acidosis Increases p-Glycoprotein Activity and Chemoresistance in Tumors in Vivo via p38 Signaling Pathway. held in Boston, MA, 2011//. J.C. LaManna, M.A. Puchowicz, K. Xu, D.K. Harrison, and D.F. Bruley, eds. (Springer US), pp. 115–122.

Vander Heiden, M.G., Cantley, L.C., and Thompson, C.B. (2009). Understanding the Warburg effect: the metabolic requirements of cell proliferation. Science 324, 1029–1033. 10.1126/science.1160809.

Vaneycken, I., Devoogdt, N., Van Gassen, N., Vincke, C., Xavier, C., Wernery, U., Muyldermans, S., Lahoutte, T., and Caveliers, V. (2011). Preclinical screening of anti-HER2 nanobodies for molecular imaging of breast cancer. Faseb j 25, 2433–2446. 10.1096/fj.10-180331.

Wagner, T.R., and Rothbauer, U. (2021). Nanobodies - Little helpers unravelling intracellular signaling. Free Radic Biol Med 176, 46–61. 10.1016/j.freeradbiomed.2021.09.005.

Webb, B.A., Chimenti, M., Jacobson, M.P., and Barber, D.L. (2011). Dysregulated pH: a perfect storm for cancer progression. Nat Rev Cancer 11, 671–677. 10.1038/nrc3110.

Weerakkody, D., Moshnikova, A., Thakur, M.S., Moshnikova, V., Daniels, J., Engelman, D.M., Andreev, O.A., and Reshetnyak, Y.K. (2013). Family of pH (low) insertion peptides for tumor targeting. Proc Natl Acad Sci U S A 110, 5834–5839. 10.1073/pnas.1303708110.

Whitfield, J.H., Zhang, W.H., Herde, M.K., Clifton, B.E., Radziejewski, J., Janovjak, H., Henneberger, C., and Jackson, C.J. (2015). Construction of a robust and sensitive arginine biosensor through ancestral protein reconstruction. Protein Sci 24, 1412–1422. 10.1002/pro.2721.

Xu, C.F., Liu, Y., Shen, S., Zhu, Y.H., and Wang, J. (2015). Targeting glucose uptake with siRNA-based nanomedicine for cancer therapy. Biomaterials 51, 1–11. 10.1016/j.biomaterials.2015.01.068.

Yang, H., Geng, Y.-H., Wang, P., Yang, H., Zhou, Y.-T., Zhang, H.-Q., He, H.-Y., Fang, W.- G., and Tian, X.-X. (2020). Extracellular ATP promotes breast cancer invasion and chemoresistance via SOX9 signaling. Oncogene 39, 5795–5810. 10.1038/s41388-020-01402-z.

Yang, L., Li, A., Lei, Q., and Zhang, Y. (2019). Tumor-intrinsic signaling pathways: key roles in the regulation of the immunosuppressive tumor microenvironment. Journal of Hematology & Oncology 12, 125. 10.1186/s13045-019-0804-8.

Yang, Y., Wolfram, J., Boom, K., Fang, X., Shen, H., and Ferrari, M. (2013). Hesperetin impairs glucose uptake and inhibits proliferation of breast cancer cells. Cell Biochem Funct 31, 374–379. 10.1002/cbf.2905.

Zhang, W.H., Herde, M.K., Mitchell, J.A., Whitfield, J.H., Wulff, A.B., Vongsouthi, V., Sanchez-Romero, I., Gulakova, P.E., Minge, D., Breithausen, B., et al. (2018). Monitoring hippocampal glycine with the computationally designed optical sensor GlyFS. Nat Chem Biol 14, 861–869. 10.1038/s41589-018-0108-2.

Zhao, Y., Araki, S., Wu, J., Teramoto, T., Chang, Y.F., Nakano, M., Abdelfattah, A.S., Fujiwara, M., Ishihara, T., Nagai, T., and Campbell, R.E. (2011). An expanded palette of genetically encoded Ca²⁺ indicators. Science 333, 1888–1891. 10.1126/science.1208592.

